# ScmR, a global regulator of gene expression, quorum sensing, pH homeostasis, and virulence in *Burkholderia thailandensis*

**DOI:** 10.1101/871871

**Authors:** Servane Le Guillouzer, Marie-Christine Groleau, Florian Mauffrey, Eric Déziel

## Abstract

The nonpathogenic soil saprophyte *Burkholderia thailandensis* is a member of the *Burkholderia pseudomallei-thailandensis-mallei* (*Bptm*) group, which also comprises the closely related human pathogens *Burkholderia pseudomallei* and *Burkholderia mallei* responsible for the diseases melioidosis and glanders, respectively. ScmR, a recently identified LysR-type transcriptional regulator (LTTR) in *B. thailandensis* acts as a global transcriptional regulator throughout the stationary phase, and modulates the production of a wide range of secondary metabolites, including *N*-acyl-L-homoserine lactones (AHLs) and 4-hydroxy-3-methyl-2-alkylquinoline (HMAQ), virulence in the model host *Caenorhabditis elegans*, as well as several quorum sensing (QS)-dependent phenotypes. We have investigated the role of ScmR in *B. thailandensis* strain E264 during the exponential phase. We used RNA-Sequencing (RNA-Seq) transcriptomic analyses to identify the ScmR regulon, which was compared to the QS-controlled regulon, showing a considerable overlap between the ScmR-regulated genes and those controlled by QS. We characterized several genes modulated by ScmR, using quantitative reverse transcription-PCR (qRT-PCR) or mini-CTX-*lux* transcriptional reporters, including the oxalate biosynthetic gene *obc1* required for pH homeostasis, the orphan LuxR-type transcriptional regulator BtaR5-encoding gene, the *bsa* (*Burkholderia* secretion apparatus) type III secretion system (T3SS) genes essential for both *B. pseudomallei* and *B. mallei* pathogenicity, as well as the *scmR* gene itself. We confirmed that the transcription of *scmR* is under QS control, presumably ensuring fine-tuned modulation of gene expression. Finally, we demonstrate that ScmR influences virulence using the fruit fly model host *Drosophila melanogaster*. We conclude that ScmR represents a central component of the *B. thailandensis* QS regulatory network.

**Importance:** Coordination of the expression of genes associated with bacterial virulence and environmental adaptation is often dependent on quorum sensing (QS). The QS circuitry of the nonpathogenic bacterium *Burkholderia thailandensis*, which is widely used as a model system for the study of the human pathogen *Burkholderia pseudomallei*, is complex. We found that the recently identified LysR-type transcriptional regulator (LTTR), ScmR, which is highly conserved and involved in the control of virulence/survival factors in the *Burkholderia* genus, is a global regulator mediating gene expression through the multiple QS systems coexisting in *B. thailandensis*, as well as independently of QS. We conclude that ScmR represents a key QS modulatory network element, ensuring tight regulation of the transcription of QS-controlled genes, particularly those required for acclimatization to the environment.

## Introduction

Quorum sensing (QS) is a global regulatory mechanism of gene expression depending on bacterial density (1). Gram-negative bacteria often possess homologues of the LuxI/LuxR system initially characterized in the bioluminescent marine bacterium *Vibrio fischeri* (2). The signaling molecules *N*-acyl-L-homoserine lactones (AHLs), which are typically produced by LuxI-type synthases, accumulate in the environment as bacterial growth progresses until a threshold concentration is reached allowing bacteria to synchronize their activities and to function as multicellular communities. These AHLs activate LuxR-type transcriptional regulators that modulate the transcription of QS target genes, which contain a *lux* box sequence in their promoter region (3).

*Burkholderia thailandensis* is a nonpathogenic soil saprophyte belonging to the *Burkholderia-pseudomallei-thailandensis-mallei* (*Bptm*) group, which also comprises the closely related pathogens *Burkholderia pseudomallei* and *Burkholderia mallei* responsible for melioidosis and glanders, respectively (4). *B. thailandensis* is considered the avirulent version of *B. pseudomallei* (5), and is thus commonly used as a surrogate model for the study of *B. pseudomallei*, which is considered a potential bioterrorism agent and whose manipulation is consequently restricted to biosafety level 3 (BSL3) laboratories (6). The members of the *Bptm* group carry multiple LuxI/LuxR QS systems that are associated with the biosynthesis of numerous AHL signaling molecules (4, 7–9). These QS systems are referred to as the BtaI1/BtaR1 (QS-1), BtaI2/BtaR2 (QS-2), and BtaI3/BtaR3 (QS-3) QS systems in *B. thailandensis* (10, 11). The QS-1 system is composed of the BtaR1 transcriptional regulator and the BtaI1 synthase, which synthesizes *N-*octanoyl-L-homoserine lactone (C_8_-HSL) (12, 13). The BtaR2 transcriptional regulator and the BtaI2 synthase that catalyze the biosynthesis of both *N-*3-hydroxy-decanoyl-L-homoserine lactone (3OHC_10_-HSL) and *N-*3-hydroxy-octanoyl-L-homoserine lactone (3OHC_8_-HSL) constitute the QS-2 system (12, 14). The QS-3 system is composed of the BtaR3 transcriptional regulator and the BtaI3 synthase is also responsible for 3OHC_8_-HSL production (12, 13). Furthermore, *B. thailandensis*, *B. pseudomallei*, and *B. mallei* carry orphan *luxR* homologues, namely, *btaR4* (*malR*) and *btaR5* in *B. thailandensis* (15, 16).

QS is involved in the regulation of several virulence factors in *B. pseudomallei* and *B. mallei*, and is essential to their full capacity to cause infections (7, 8, 17, 18). Other QS-controlled phenotypic traits among the *Bptm* group members have been reported, such as colony morphology, the development of biofilm, self-aggregation, motility, pH homeostasis, as well as production of secondary metabolites (9, 10, 13, 14, 16, 18–25).

A LysR-type transcriptional regulator (LTTR) involved in secondary metabolism regulation, hence designated ScmR, was recently identified in the *Bptm* group members (26). LTTRs are part of a large family and display a well conserved structure with a N-terminal DNA-binding helix-turn-helix motif and a C-terminal cofactor-binding domain (27). LTTRs are typically negatively autoregulated and frequently positively modulate expression of adjacent genes (27). Nevertheless, LTTRs were also described as global regulators acting positively or negatively (27). Mao *et al.* (26) demonstrated that ScmR constitutes a global regulator of gene expression in *B. thailandensis* and influences the production of a wide range of secondary metabolites, including AHLs and the putative 4-hydroxy-3-methyl-2-alkylquinoline (HMAQ) signaling molecules, virulence in the nematode worm model *Caenorhabditis elegans*, as well as several QS-dependent phenotypes. Additionally, expression of the *scmR* gene is under QS control (10, 26).

The central goal of the present study was to further characterize the molecular mechanism of action of the *B. thailandensis* E264 ScmR transcriptional regulator. We found that ScmR is a global regulator mediating gene expression through the QS-1, QS-2, and/or QS-3 systems, as well as independently of QS. Furthermore, we identified novel genes modulated by ScmR, including the oxalate biosynthetic gene *obc1* that is essential for pH homeostasis in the *Burkholderia* genus, the orphan LuxR-type transcriptional regulator BtaR5-encoding gene, and the *bsa* (*Burkholderia* secretion apparatus) type III secretion system (TTSS) genes required for both *B. pseudomallei* and *B. mallei* pathogenicity. Moreover, we showed that *scmR* is negatively autoregulated, and we confirmed that its transcription is QS-controlled, ensuring tight regulation of gene expression by ScmR in *B. thailandensis*. Finally, we demonstrate that ScmR represses virulence using the fruit fly model *Drosophila melanogaster*. All in all, this study contributes to a better appreciation of the ScmR regulatory mechanism of the expression of genes in *B. thailandensis*, and in particular those related to virulence of *B. pseudomallei*.

## Results

### The ScmR regulon comprises many QS-controlled genes

ScmR was recently described as a global transcriptional regulator impacting gene expression during the stationary phase of bacterial growth in *B. thailandensis* (26). We used RNA-Seq transcriptomic analyses to further characterize the regulon of the ScmR transcriptional regulator. We identified the ScmR-regulated genes by comparing the transcripts in the wild-type and in the *scmR*-mutant strains of *B. thailandensis* E264 throughout the logarithmic growth phase. We found that ScmR both positively and negatively influence the expression of genes located on the two *B. thailandensis* E264 chromosomes (Fig. 1A). Using a 3-fold difference in transcription as a cut-off, we identified 907 genes that were positively modulated by ScmR, and 397 genes that were negatively modulated by ScmR (Fig. 1A). These findings confirm that ScmR constitutes a global regulator of gene expression in *B. thailandensis* E264 (26). Our RNA-Seq analyses identified genes known to be controlled by ScmR or genes encoding functions known to be controlled by ScmR. Indeed, Mao *et al.* (26) recently demonstrated that ScmR stimulates the production of HMAQ, which includes putative signals. RNA-Seq confirmed that expression of the *hmqABCDEFG* operon, which is required for HMAQs production (28), is activated by ScmR (**Table S1**). Furthermore, ScmR represses the production of burkholdac, a hybrid polyketide/nonribosomal peptide and a potent inhibitor of some histone deacetylases (HDACs) (26). Consistently, expression of the *bhc* gene cluster, responsible for burkholdac biosynthesis (29), was increased in the *scmR*-mutant compared to the wild-type strain (**Table S1**). Moreover, we observed that ATP synthesis and stress response genes were downregulated in the absence of ScmR (**Table S1**), as recently reported (26). Finally, RNA-Seq showed that transcription of the putative exopolysaccharide (EPS) genes *bceABCDEFGHIJ* and *bceNOPRSTU* is affected by ScmR (**Table S1**). This is in agreement with the finding that ScmR influences colony morphology, as well as pellicle and biofilm formation of *B. thailandensis* E264 (26).

**Figure 1.**
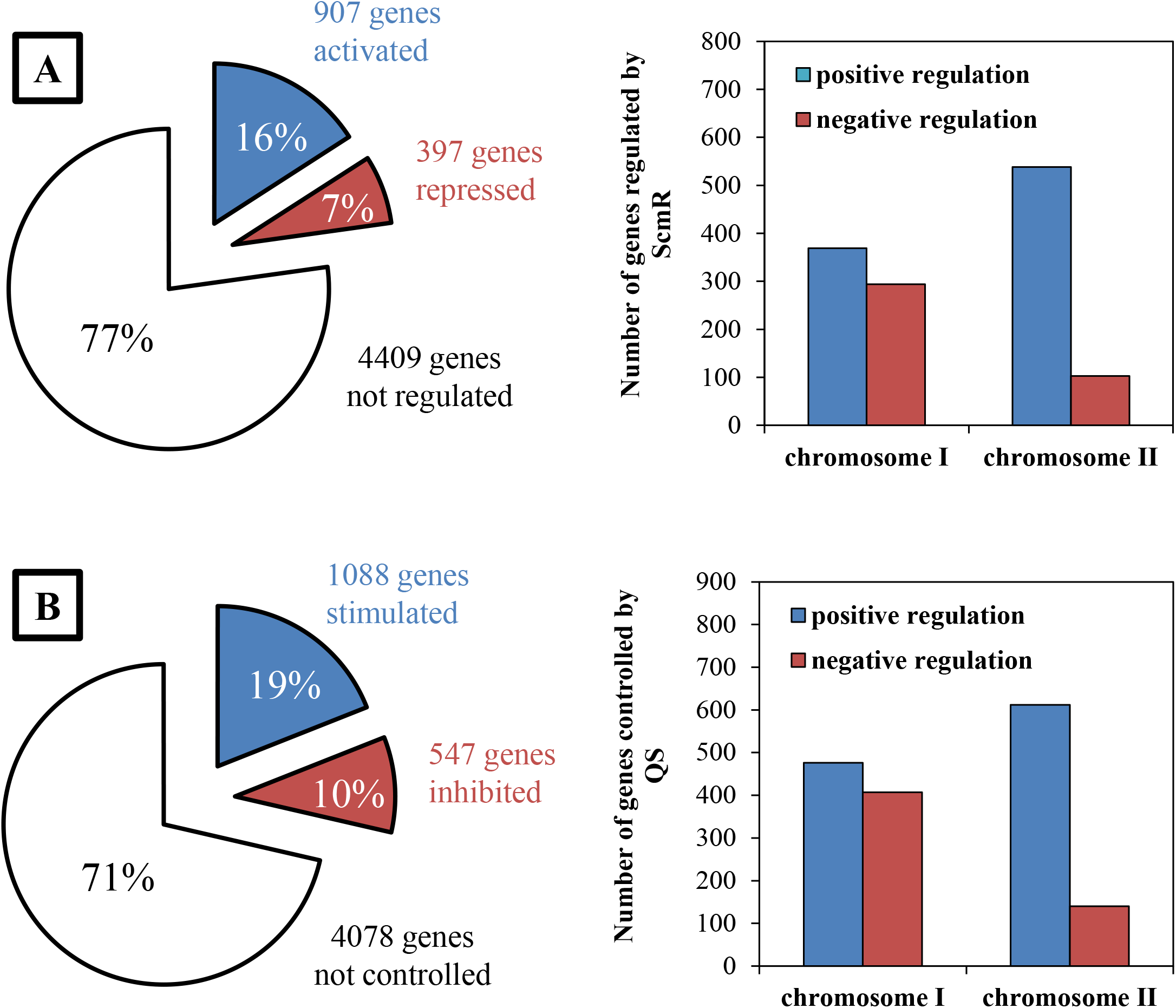
Numbers of ScmR-regulated genes and those QS-controlled in *B. thailandensis* E264 according to transcriptomic analyses obtained by RNA-Seq. (A) Total numbers of genes showing positive or negative modulation in the *scmR*-mutant compared to the wild-type strain of *B. thailandensis* E264. (B) Total numbers of genes showing positive or negative modulation in the Δ*btaI1*Δ*btaI2*Δ*btaI3* mutant in comparison with the wild-type strain of *B. thailandensis* E264.

The ScmR transcriptional regulator was shown to influence the biosynthesis of C_8_-HSL, 3OHC_10_-HSL, and 3OHC_8_-HSL (26), the main AHLs produced by *B. thailandensis* E264 (10, 11, 13, 14). Therefore, we assumed that ScmR could intervene in the regulation of gene expression, *inter alia*, by impacting the QS-1, QS-2, and/or QS-3 systems of *B. thailandensis* E264. Indeed, Mao *et al.* (26) demonstrated that QS-dependent phenotypes, including colony morphology, as well as the development of biofilm, are influenced by ScmR and we accordingly found several previously reported QS-controlled genes in the ScmR regulon (**Table S1**). Consequently, we also compared the transcripts in the wild-type strain of *B. thailandensis* E264 and in the AHL-defective Δ*btaI1*Δ*btaI2*Δ*btaI3* mutant under the same growth conditions to identify the genes specifically modulated by ScmR independently of its effect on QS. Our RNA-Seq analyses indicate that QS positively regulated expression of 1088 genes and negatively modulated expression of 547 genes on both chromosomes of *B. thailandensis* E264 (Fig. 1B). Importantly, we confirmed the involvement of QS in the regulation of genes affected by AHLs or genes encoding functions affected by AHLs. In *B. thailandensis*, QS stimulates contact-dependent growth inhibition (CDI) (10, 30), and we indeed observed that the transcription of the CDI genes was decreased in the absence of AHLs (**Table S1**). Furthermore, RNA-Seq indicated that the transcription of the bactobolin biosynthetic genes (14), as well as the *obc1* gene expression, encoding the oxalate biosynthetic enzyme Obc1 that is essential to pH homeostasis (19), are activated by QS (**Table S1**), as previously reported (10). Moreover, RNA-Seq confirmed that expression of both flagellar genes and methyl-accepting chemotaxis protein genes was upregulated in the Δ*btaI1*Δ*btaI2*Δ*btaI3* mutant in comparison with the wild-type strain (10) (**Table S1**), which is consistent with the observation that *B. thailandensis* E264 QS mutants are hypermotile (13).

Interestingly, we found a considerable overlap between the genes regulated by ScmR and those QS-controlled (**Table S2**). We identified 681 genes activated by both ScmR and QS (Fig. 2A), whereas 310 genes were repressed by both ScmR and QS (Fig. 2B). Other patterns of coregulation were observed including positive regulation by ScmR and negative regulation by QS (Fig. 2C), as well as negative regulation by ScmR and positive regulation by QS (Fig. 2D). While we identified 1019 genes that were coregulated by both ScmR and QS, 901 genes appeared to be independently regulated by either ScmR or QS under the conditions of our experiments (Fig. 2). Altogether, these results support the idea that ScmR regulates the transcription of many genes through modulation of the QS-1, QS-2 and/or QS-3 systems in *B. thailandensis* E264. Additionally, we found that ScmR affected the expression of genes encoding transcriptional factors, including the QS-controlled orphan transcriptional regulator BtaR5-encoding gene (**Table S2**). Thus, many genes could be modulated by ScmR indirectly through auxiliary regulators, as recently proposed (26).

**Figure 2.**
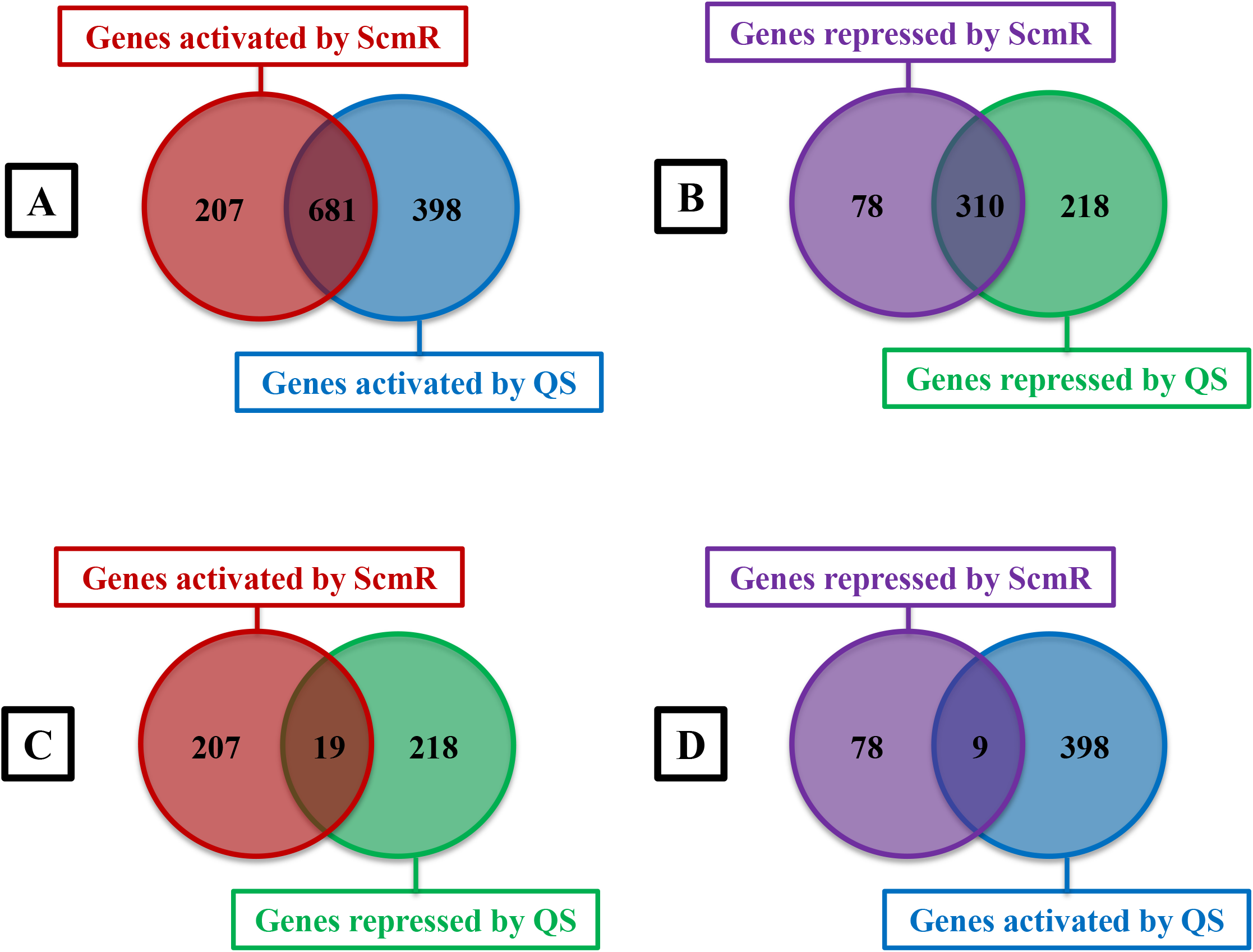
The ScmR regulon comprises many QS-controlled genes. (A) Number of genes positively regulated by ScmR and/or activated by QS. (B) Number of genes negatively modulated by ScmR and/or repressed by QS. (C) Number of genes stimulated by ScmR and/or negatively controlled by QS. (D) Number of genes inhibited by ScmR and/or positively controlled by QS.

### ScmR modulates AHLs biosynthesis but not the transcription of the AHL synthase-coding genes

The influence of ScmR on C_8_-HSL, 3OHC_10_-HSL, and 3OHC_8_-HSL production was demonstrated throughout the stationary phase of growth (26), but its effect on QS during the logarithmic phase had not been investigated yet. To determine whether the biosynthesis of the main AHLs produced by *B. thailandensis* E264 was impacted by ScmR in the exponential phase, we compared the concentrations of these AHLs in the wild-type strain of *B. thailandensis* E264 and in the *scmR*-mutant. We confirmed that the levels of C_8_-HSL were decreased in the absence of ScmR (Fig. 3A), as previously reported (26), suggesting that ScmR is an activator on the QS-1 system. In contrast to stationary phase observations (26), however, we detected increasing concentrations of 3OHC_10_-HSL and 3OHC_8_-HSL in the *scmR*-mutant versus the wild-type strain under our conditions (Figs. 3B and C), indicating that the QS-2 and/or QS-3 system might be repressed by ScmR.

**Figure 3.**
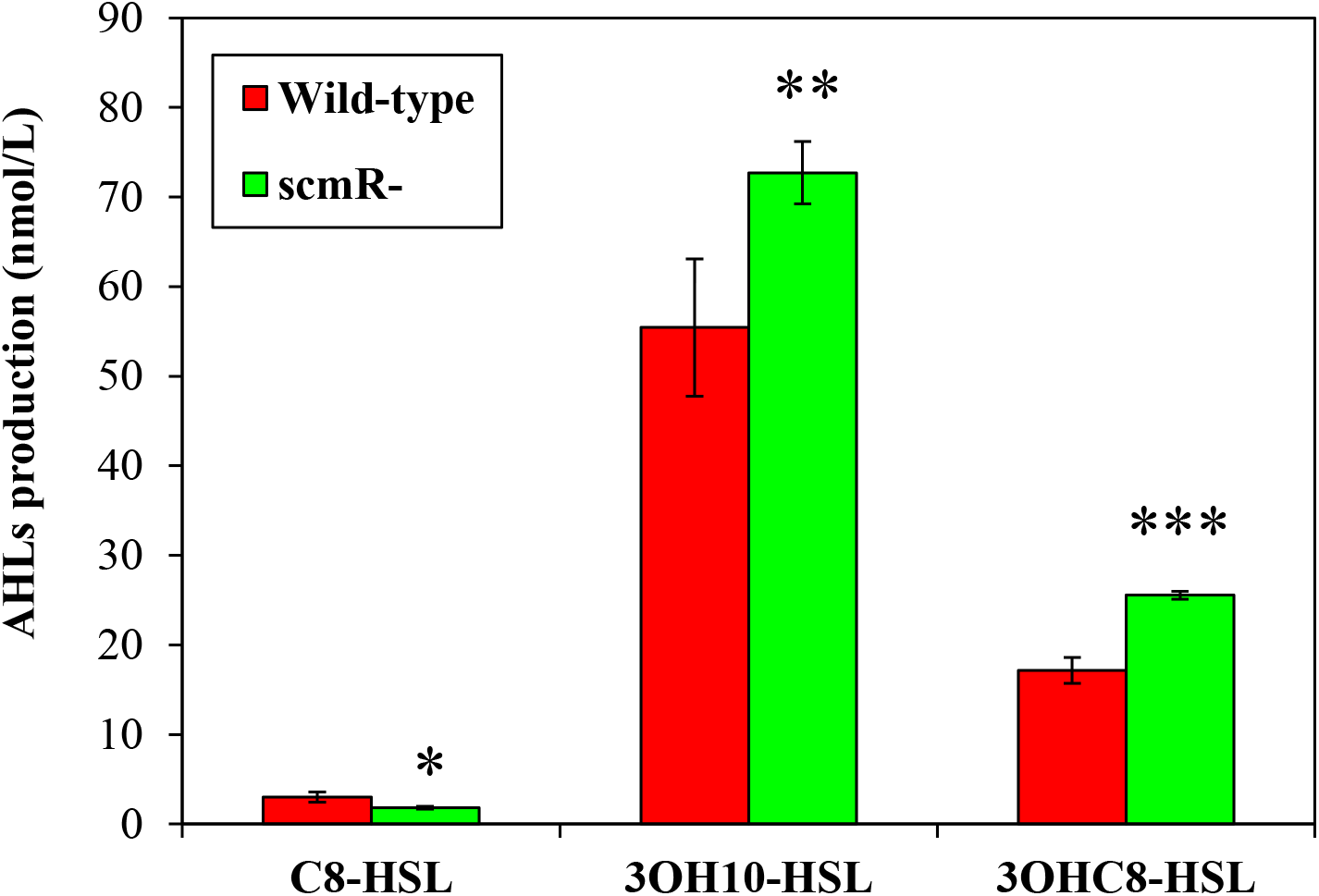
ScmR influences the production of the main AHLs found in *B. thailandensis* E264 during the logarithmic phase of growth. C_8_-HSL, 3OHC_10_-HSL, and 3OHC_8_-HSL biosynthesis was measured using LC-MS/MS in cultures of the wild-type and of the *scmR*-mutant strains of *B. thailandensis* E264. The values are means ± standard deviations (error bars) for three replicates. Values that are significantly different are indicated by asteriks as follows: ***, *P* < 0.001; **, *P* < 0.01; *, *P* < 0.05.

ScmR stimulates the production of C_8_-HSL, 3OHC_10_-HSL, and 3OHC_8_-HSL during the stationary phase, but the expression of the BtaI1-, BtaI2-, and BtaI3-encoding genes responsible for the production of these AHLs, nor the transcription of *btaR1*, *btaR2*, and *btaR3* were downregulated in an Δ*scmR* mutant in comparison with the wild-type (26). To gain insights into the ScmR-dependent modulation of C_8_-HSL, 3OHC_10_-HSL, and 3OHC_8_-HSL biosynthesis, we determined the expression profiles of the AHL synthase-coding genes *btaI1*, *btaI2*, and *btaI3* throughout the bacterial growth phases in cultures of the *scmR*-mutant versus the *B. thailandensis* E264 wild-type strain harboring a chromosomal *btaI1*-*lux*, *btaI2*-*lux*, or *btaI3*-*lux* transcriptional fusion. No discernible difference in expression from the *btaI1*, *btaI2*, and *btaI3* promoters was found in the *scmR*-mutant compared to the wild-type strain (data not shown). Accordingly, our RNA-Seq analyses indicated that ScmR had no impact on *btaI1*, *btaI2*, and *btaI3* transcription (**Table S1**). The *btaR1*, *btaR2*, and *btaR3* genes, encoding the BtaR1, BtaR2, and BtaR3 transcriptional regulators, respectively, were not affected by ScmR neither (**Table S1**). Taken together, these data confirm that the effect of ScmR on C_8_-HSL, 3OHC_10_-HSL, and 3OHC_8_-HSL biosynthesis does not result from modulation of expression of the QS-1, QS-2, and/or QS-3 system genes.

### ScmR contributes to pH homeostasis

Interestingly, we noticed growth differences between the *B. thailandensis* E264 wild-type strain and the *scmR*-mutant under the conditions of our experiments. Indeed, inactivation of *scmR* results in reduced OD_600_ during the stationary phase, but not during the exponential phase (Fig. 4). Since pH was reported to significantly affect the growth of *B. thailandensis* E264, *B. pseudomallei* Bp82, and *Burkholderia glumae* BGR1 (19, 31), we hypothesized that it could be involved in the *scmR-* mutant phenotype. We analyzed the implication of ScmR in pH homeostasis by measuring the pH in cultures of the *B. thailandensis* E264 wild-type strain and the *scmR*-mutant throughout the different stages of the bacterial growth. pH in cultures of both the wild-type strain and the *scmR*-mutant was approximately 7.3 during the exponential phase (Fig. 5A). On the other hand, pH in wild-type strain cultures decreased to between 7.0 and 6.5 throughout the stationary phase, whereas pH in *scmR*-mutant cultures increased to between 7.5 and 8.0, apparently correlating with the OD_600_ stabilization (Figs. 5B and C). To verify whether growth inhibition could be caused by alkaline toxicity, we buffered cultures of the *scmR*-mutant with 100 mM HEPES (pH 7.0) and observed that the effect on the OD_600_ was alleviated (Figs. 4 and **5**), supporting the hypothesis that culture medium alkalization is the cause of the *scmR*-mutant growth differences.

**Figure 4.**
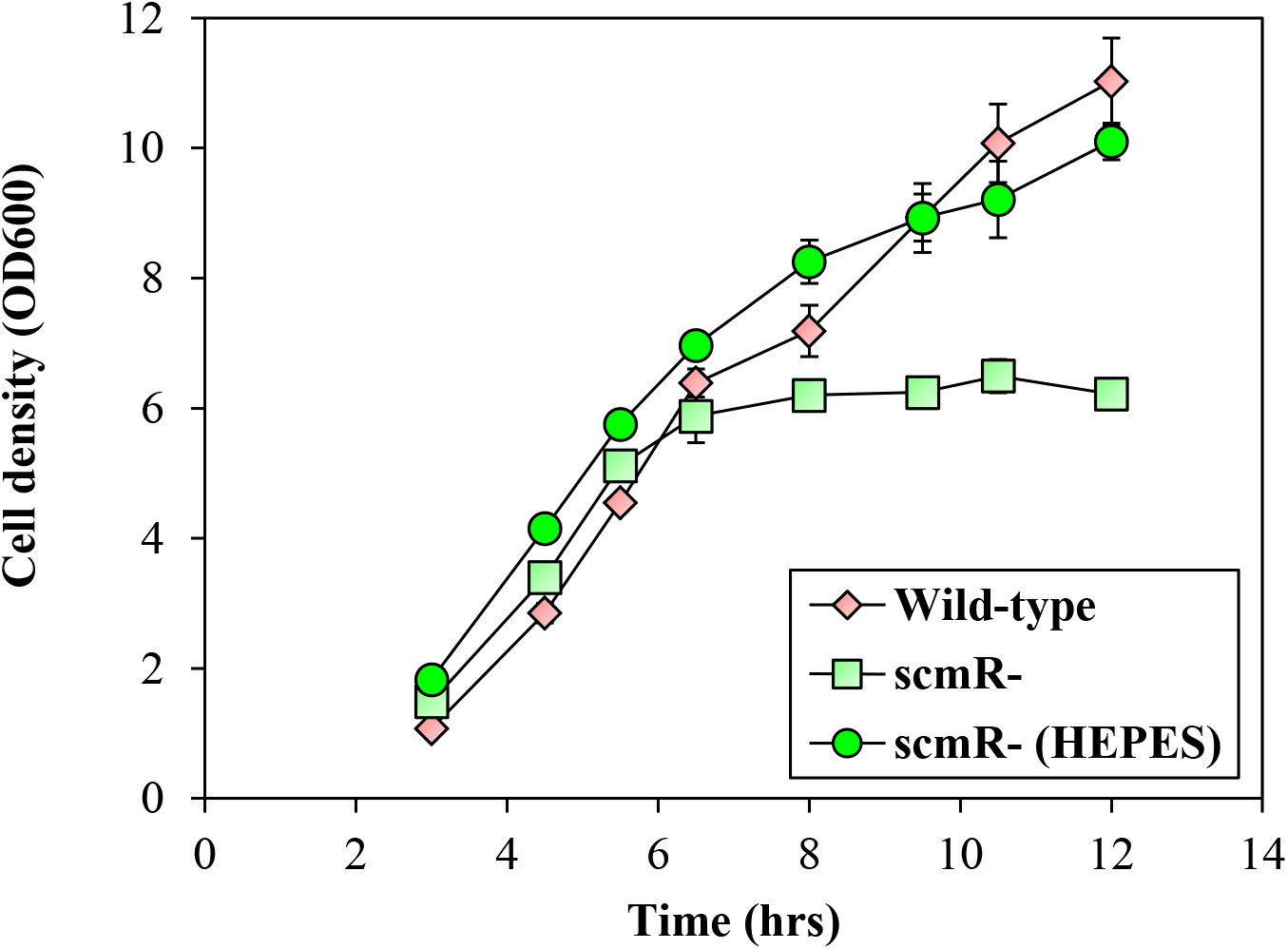
Impact of *scmR* inactivation on bacterial growth. The *B. thailandensis* E264 wild-type strain and the *scmR*-mutant strain growth curves. Cultures were buffered with 100 mM HEPES. Water only was added to the controls. The error bars represent the standard deviations of the averages for three replicates.

**Figure 5.**
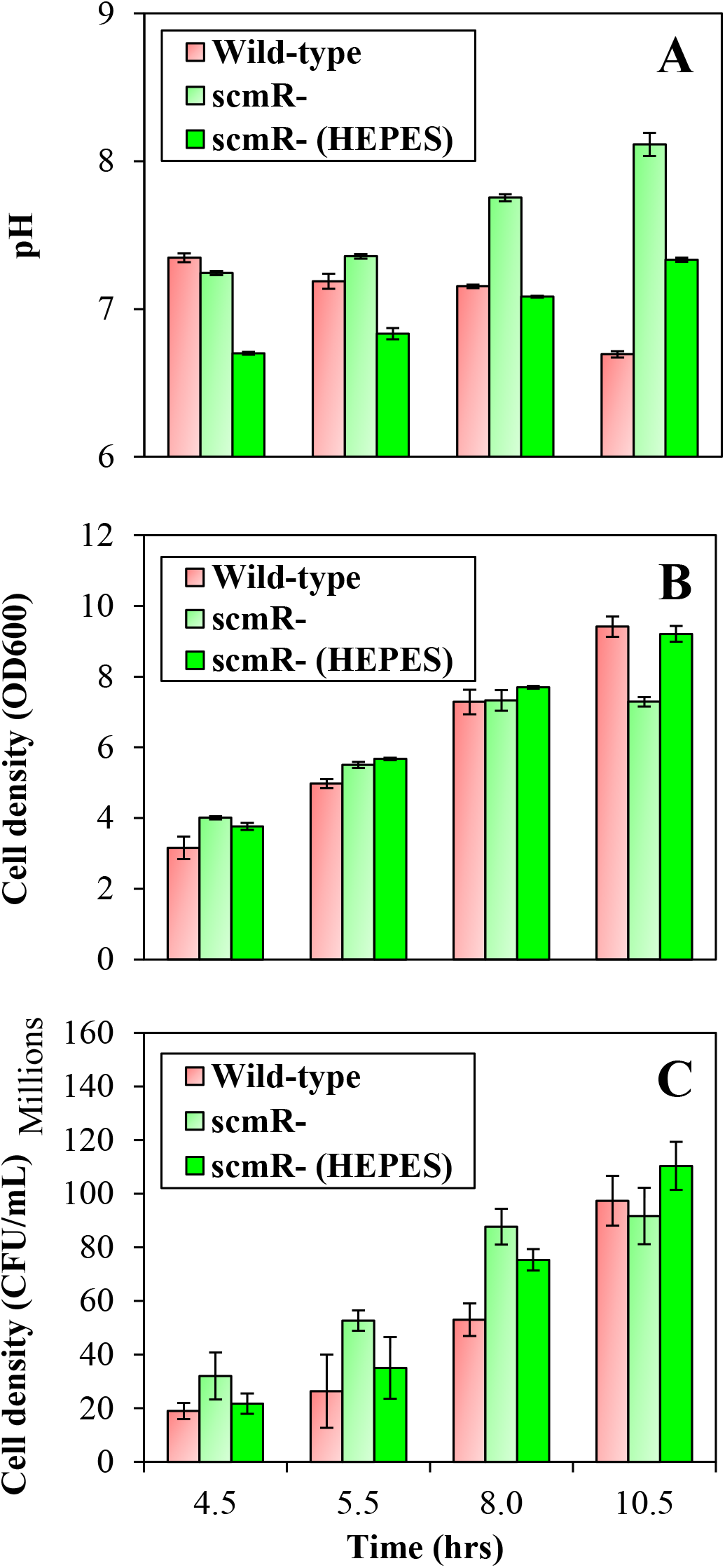
ScmR influences pH homeostasis. (A) pH value was measured with a pH electrode and meter (Mettler-Toledo, Mississauga, ON, Canada), at various times during growth in cultures of the *B. thailandensis* E264 wild-type strain and the *scmR*-mutant strain. Cultures were buffered with 100 mM HEPES. Water only was added to the controls. (B) The cell density was monitored by measuring turbidity, expressed in 600 nm absorption units (OD_600_). (C) Colony-forming units (CFUs) were determined by plate-counting methods. The error bars represent the standard deviations of the averages for three replicates.

To further characterize the underlying regulatory mechanisms directing pH homeostasis through ScmR, we investigated the effect of ScmR on expression of the *obc1* gene, encoding the oxalate biosynthetic enzyme Obc1, which influences the pH in several *Burkholderia* spp. (19, 31). Oxalic acid was indeed reported to be essential to neutralize alkalization in stationary-phase cultures of the wild-type strain of *B. thailandensis* E264, *B. pseudomallei* Bp82, and *B. glumae* BGR1 (19, 31). Expression of *obc1*, as well as oxalic acid production, were both shown to be QS-controlled (10, 19). Our RNA-Seq analyses indicate that *obc1* transcription was downregulated in the AHL-defective Δ*btaI1*Δ*btaI2*Δ*btaI3* mutant in comparison with the wild-type strain (approximately 35-fold) (**Table S2**), confirming that QS activates the *obc1* gene expression. Furthermore, we noticed a drastic reduction in transcription of *obc1* in the *scmR*-mutant compared to the wild-type strain (approximately 72-fold) (**Table S2**), revealing that the transcription of *obc1* is also strongly enhanced by ScmR. To ascertain the involvement of ScmR in the regulation of the *obc1* gene, expression of *obc1* was assessed by qRT-PCR in cultures of the *B. thailandensis* E264 wild-type strain and the *scmR*-mutant buffered or not with HEPES during the logarithmic growth phase. We observed that the transcription of *obc1* was completely abolished in the absence of ScmR (Fig. 6A), attesting that expression of *obc1* is tightly controlled by ScmR. Moreover, the finding that *obc1* expression is stimulated by ScmR under neutral conditions confirms previous observations that alkaline stress does not induce *obc1* transcription (10, 19). All in all, these findings suggest that ScmR intervenes in pH homeostasis by regulating oxalic acid biosynthesis.

**Figure 6.**
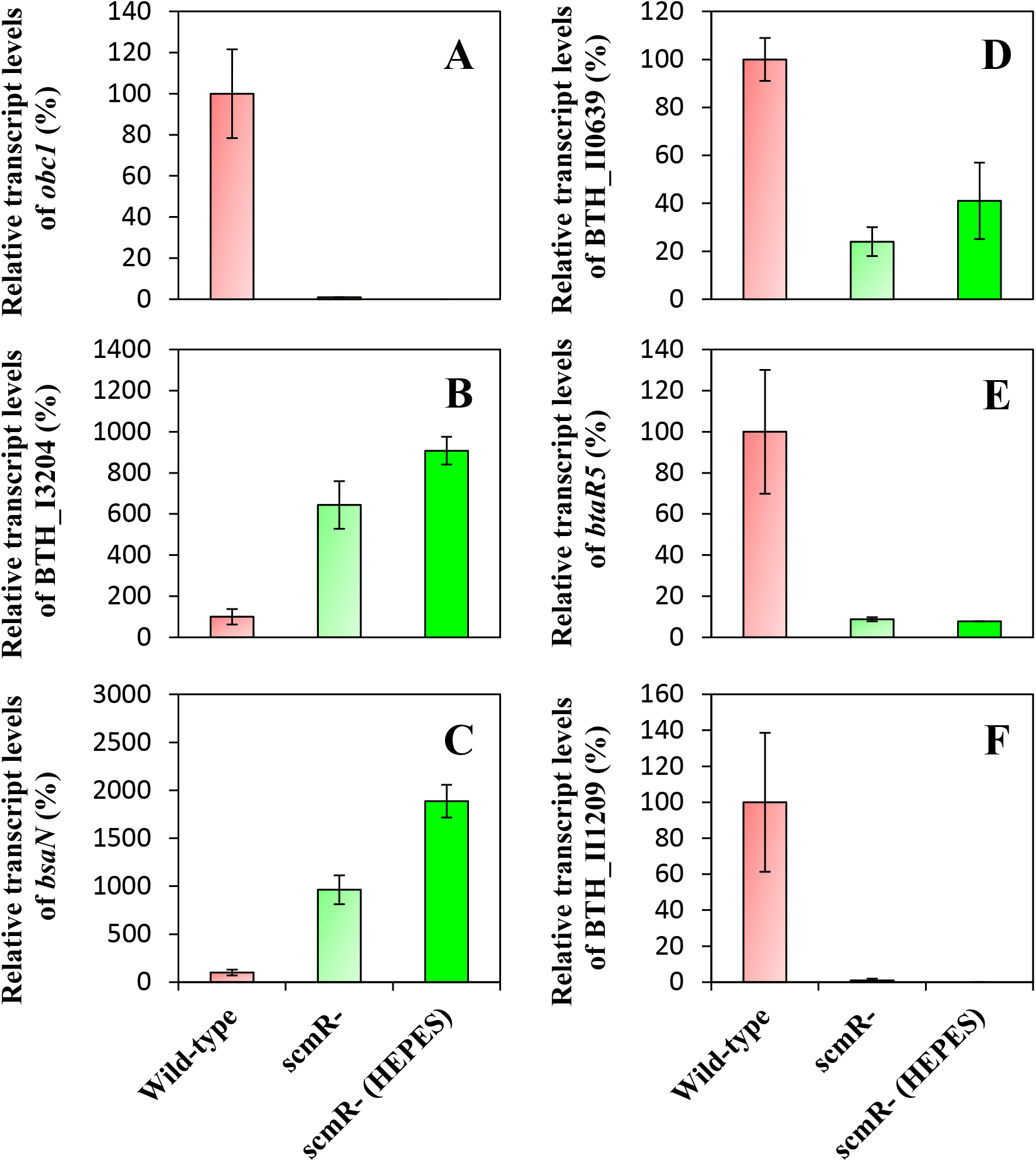
Expression of several ScmR-regulated genes in cultures buffered or not with HEPES of the *B. thailandensis* E264 wild-type and the *scmR-* mutant strains. The relative transcript levels of (A) *obc1*, (B) BTH_I3204, (C) *bsaN*, (D) BTH_II0639, (E) *btaR5*, and (F) BTH_II1209 were assessed by qRT-PCR in cultures of the wild-type and of the *scmR-* mutant strains of *B. thailandensis* E264. Cultures were buffered with 100 mM HEPES. Water only was added to the controls. The results are presented as relative quantification of transcription of the gene compared to the wild-type strain, which was set at 100%. The error bars represent the standard deviations of the averages for three replicates.

Considering the impact of the absence of ScmR on pH, we hypothesized that some of the regulatory effects observed in our RNA-Seq analyses could result from pH imbalance. Consequently, we measured the transcription of several genes that were affected in the *scmR*-mutant strain in comparison with the wild-type strain of *B. thailandensis* E264, namely, BTH_I3204, *bsaN*, BTH_II0639, *btaR5*, and BTH_II1209 encoding a lipoprotein, the T3SS transcriptional regulator BsaN, a lipase, the orphan LuxR-type transcriptional regulator BtaR5, and a hypothetical protein, respectively. Their expression was monitored by qRT-PCR during the exponential phase in cultures of the *B. thailandensis* E264 wild-type and the *scmR*-mutant strains supplemented or not with HEPES. According to our transcriptomic data, expression of the BTH_I3204 and *bsaN* genes is repressed by ScmR (approximately 26-fold and 17-fold, respectively), whereas expression of the BTH_II0639, *btaR5*, and BTH_II1209 genes is stimulated by ScmR (approximately 16-fold, 27-fold, and 274-fold, respectively) (**Table S1**). We observed that buffering *scmR-* mutant cultures did not restore normal expression of any of these genes to wild-type levels, showing that the effects observed on these genes in the *scmR-* mutant does not result from culture medium alkalization.

pH affects the integrity of AHL signaling molecules. AHLs are stable at neutral and acidic pH, while alkaline conditions cause AHLs hydrolysis (32, 33). Therefore, we asked whether ScmR could influence C_8_-HSL, 3OHC_10_-HSL, and 3OHC_8_-HSL stability by impacting pH homeostasis. Concentrations of C_8_-HSL, 3OHC_10_-HSL, and 3OHC_8_-HSL were monitored in the *B. thailandensis* E264 wild-type strain and the *scmR*-mutant throughout the different stages of bacterial growth. We confirmed that the levels of C_8_-HSL were reduced in the *scmR*-mutant in comparison with the wild-type strain in the early stages of the bacterial growth (Fig. 7A), whereas 3OHC_10_-HSL and 3OHC_8_-HSL concentrations were increased (Figs. 7B and C). As expected, the concentrations of all three AHLs were decreased in the *scmR*-mutant cultures in the late stages of bacterial growth (Fig. 7). We then examined the effect of pH buffering on AHLs levels in *scmR*-mutant cultures. The production of all three AHLs was increased in buffered cultures of the *scmR*-mutant (Fig. 7). Taken together, these observations indicate that the impact of ScmR on the QS-1, QS-2, and/or QS-3 systems might result, *inter alia*, from its influence on pH homeostasis.

**Figure 7.**
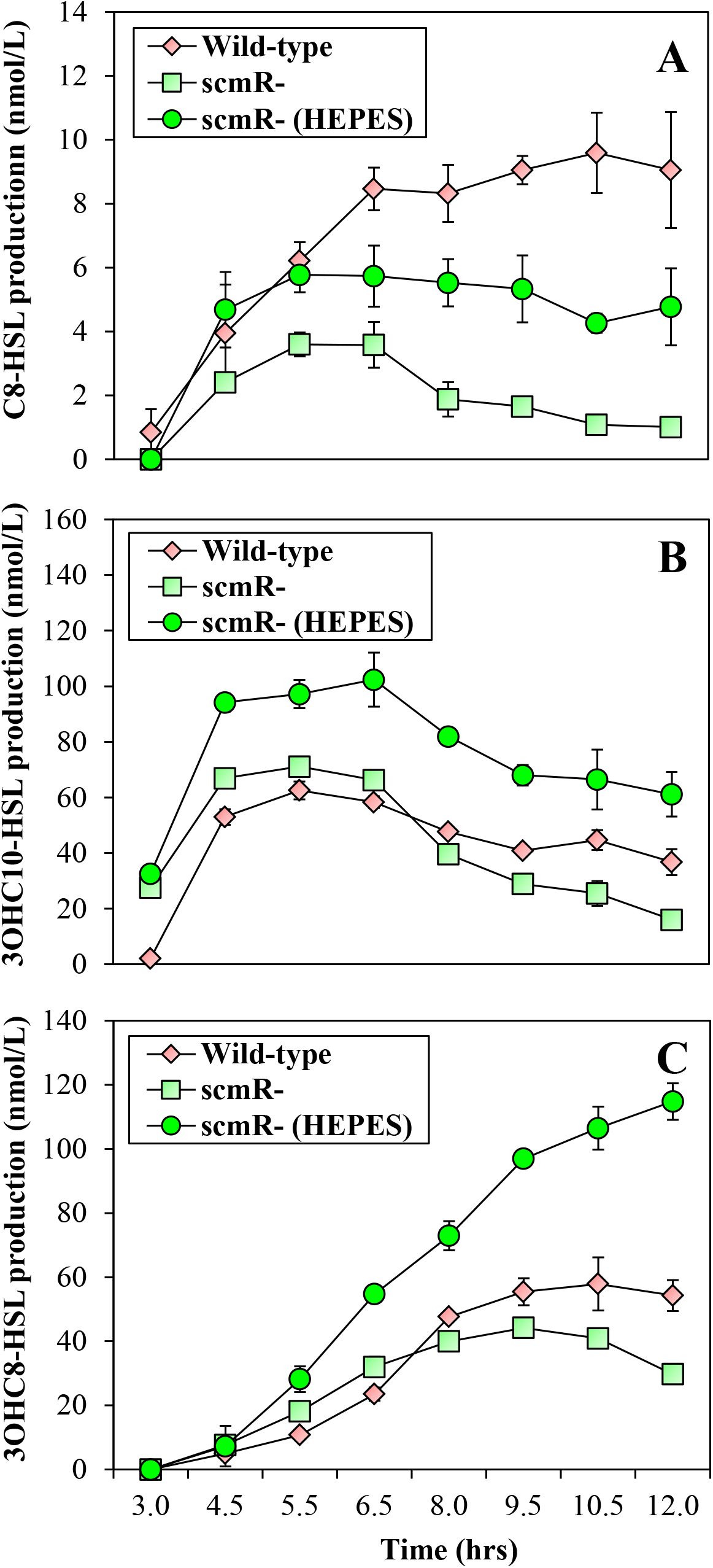
Effect of culture medium alkalization on the levels of AHLs in *B. thailandensis* E264 cultures. The concentrations of (A) C_8_-HSL, (B) 3OHC_10_-HSL, and (C) 3OHC_8_-HSL was assessed using LC-MS/MS throughout the bacterial growth phases in cultures of the wild-type and of the *scmR*-mutant strains of *B. thailandensis* E264. Cultures were buffered with 100 mM HEPES. Water only was added to the controls. The error bars represent the standard deviations of the averages for three replicates.

### QS regulation of *scmR* gene

Transcription of the *scmR* gene is activated by QS (10, 26). It was established that all three AHLs stimulate *scmR* expression (10). Accordingly, our RNA-Seq analyses indicate that *scmR* transcription is diminished in the AHL-null Δ*btaI1*Δ*btaI2*Δ*btaI3* mutant compared to the wild-type strain (approximately 4-fold) (**Table S1**), confirming that expression of *scmR* is positively modulated by QS. However, respective influence of the BtaR1, BtaR2, and BtaR3 regulators on *scmR* expression was not investigated (10). To gain insights into the QS-dependent modulation of the *scmR* gene, we measured its transcription in the Δ*btaR1*, Δ*btaR2*, and Δ*btaR3* mutants versus the *B. thailandensis* E264 wild-type strain during the logarithmic growth phase. While no obvious change in *scmR* transcription was visible in the absence of the BtaR2 transcriptional regulator, expression of *scmR* was decreased in both the Δ*btaR1* and Δ*btaR3* mutants (Fig. 8). Collectively, these observations indicate that the transcription of *scmR* is stimulated by the QS-1 and QS-3 systems, whereas the QS-2 system is not apparently involved in the modulation of *scmR* expression.

**Figure 8.**
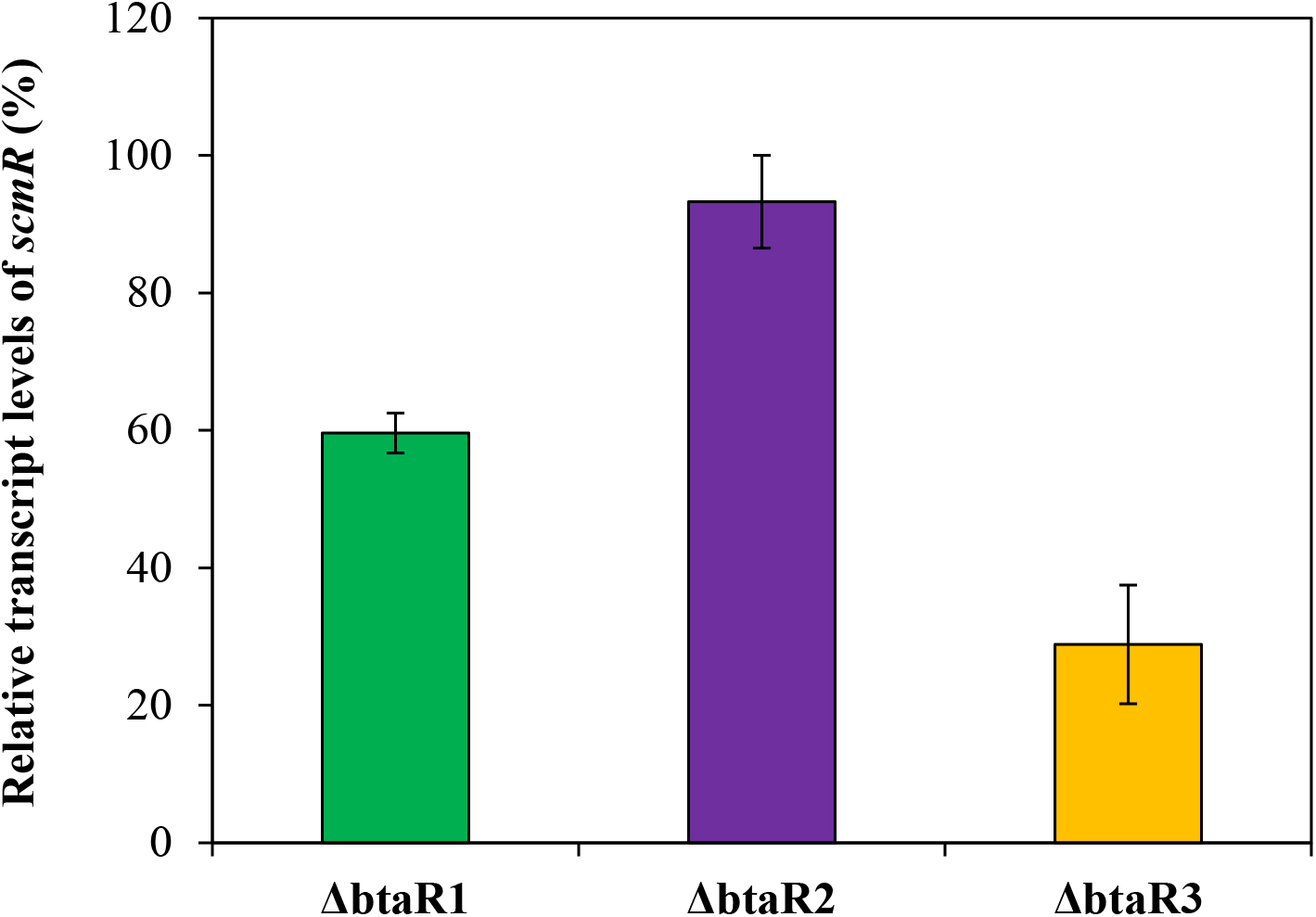
The QS-1 and QS-3 systems activate the transcription of *scmR*. The relative transcript levels of *scmR* were assessed by qRT-PCR in cultures of the wild-type and of the Δ*btaR1*, Δ*btaR2*, and Δ*btaR3* mutant strains of *B. thailandensis* E264. The results are presented as relative quantification of transcription of the gene compared to the wild-type strain, which was set at 100%. The error bars represent the standard deviations of the averages for three replicates.

While a putative *lux* box sequence was found in the promoter region of the *B. thailandensis* E264 *scmR* gene (26), we do not know whether the BtaR1 and/or BtaR3 transcriptional regulators directly control its transcription. We found a putative *lux* box sequence in the promoter region of *scmR* homologs in both *B. pseudomallei* K96243 and *B. mallei* ATCC 23344 (**Fig. S1**). Accordingly, Klaus *et al.* (34) and Majerczyk *et al.* (20) demonstrated that the expression of *scmR* in *B. pseudomallei* Bp82 and *B. mallei* GB8 is stimulated by QS, respectively. *Burkholderia cenocepacia* J2315 also possesses an *scmR* homologue, which was shown to be QS-controlled in *B. cenocepacia* K56-2 (35), but no putative *lux* box sequence was found in its promoter region (36). Altogether, these observations suggest that the QS-dependent regulation of the *scmR* gene is conserved among *Burkholderia* spp.

Since *scmR* is directly adjacent to its downstream gene, namely, *ldhA*, encoding a putative lactate dehydrogenase, on the genome of *B. thailandensis* E264, *B. pseudomallei* K96243, *B. mallei* ATCC 2344, and *B. cenocepacia* J2315, and transcribed in the same direction (**Fig. S2A**), we asked whether they could be cotranscribed. The *scmR* gene is indeed predicted to be arranged in operon with *ldhA* (http://www.burkholderia.com/), and we observed that *ldhA* transcription is also activated by QS (**Table S2**). However, both our transcriptomic data (**Fig. S2B**) and RT-PCR experiments (**Fig. S2C**) indicate that *scmR* is not cotranscribed with *ldhA*.

Interestingly, expression of *ldhA* was decreased in the *scmR*-mutant compared to the wild-type strain (**Table S2**), highlighting that the *ldhA* gene is positively modulated by ScmR as well. Of note, the reduction in expression of *ldhA* was substantially greater in the *scmR*-mutant (approximately 17-fold) than in the Δ*btaI1*Δ*btaI2*Δ*btaI3* mutant (approximately 3-fold) (**Table S2**), suggesting that QS might activate *ldhA* transcription indirectly via positive regulation of the *scmR* gene.

Since LdhA was hypothesized to influence pH homeostasis in *B. thailandensis* E264 (26), we tested its involvement in the ScmR-dependent control of pH homeostasis by measuring the pH in cultures of the *B. thailandensis* E264 wild-type strain and the *scmR-* and *ldhA*-mutants during the stationary phase of growth. While the pH in both the wild-type strain and the *ldhA*-mutant was between 6.5 and 7.0, pH in cultures of the *scmR-* mutant was approximately 9.0 (**Fig. S3A**), showing that LdhA does not affect pH in *B. thailandensis* E264 under the conditions of our experiments. Of note, inactivation of the *ldhA* gene was not associated with a change in OD_600_ (**Figs. S3B** and **S3C**). Altogether, these observations indicate that LdhA is not likely involved in the ScmR-dependent control of pH homeostasis in *B. thailandensis* E264.

### *scmR* is negatively autoregulated

As LTTRs are typically negatively autoregulated (27), we investigated the impact of ScmR on its own transcription. Considering that the use of an *scmR*-mutant to perform our RNA-Seq analyses precludes clear assessment, we measured expression of *scmR* in the *B. thailandensis* E264 wild-type strain and its *scmR*-mutant strain harboring a chromosomal *scmR*-*lux* transcriptional fusion. We observed an increase in *scmR* expression in the *scmR*-mutant in comparison with the wild-type strain (Fig. 9), revealing that *scmR* is negatively autoregulated.

**Figure 9.**
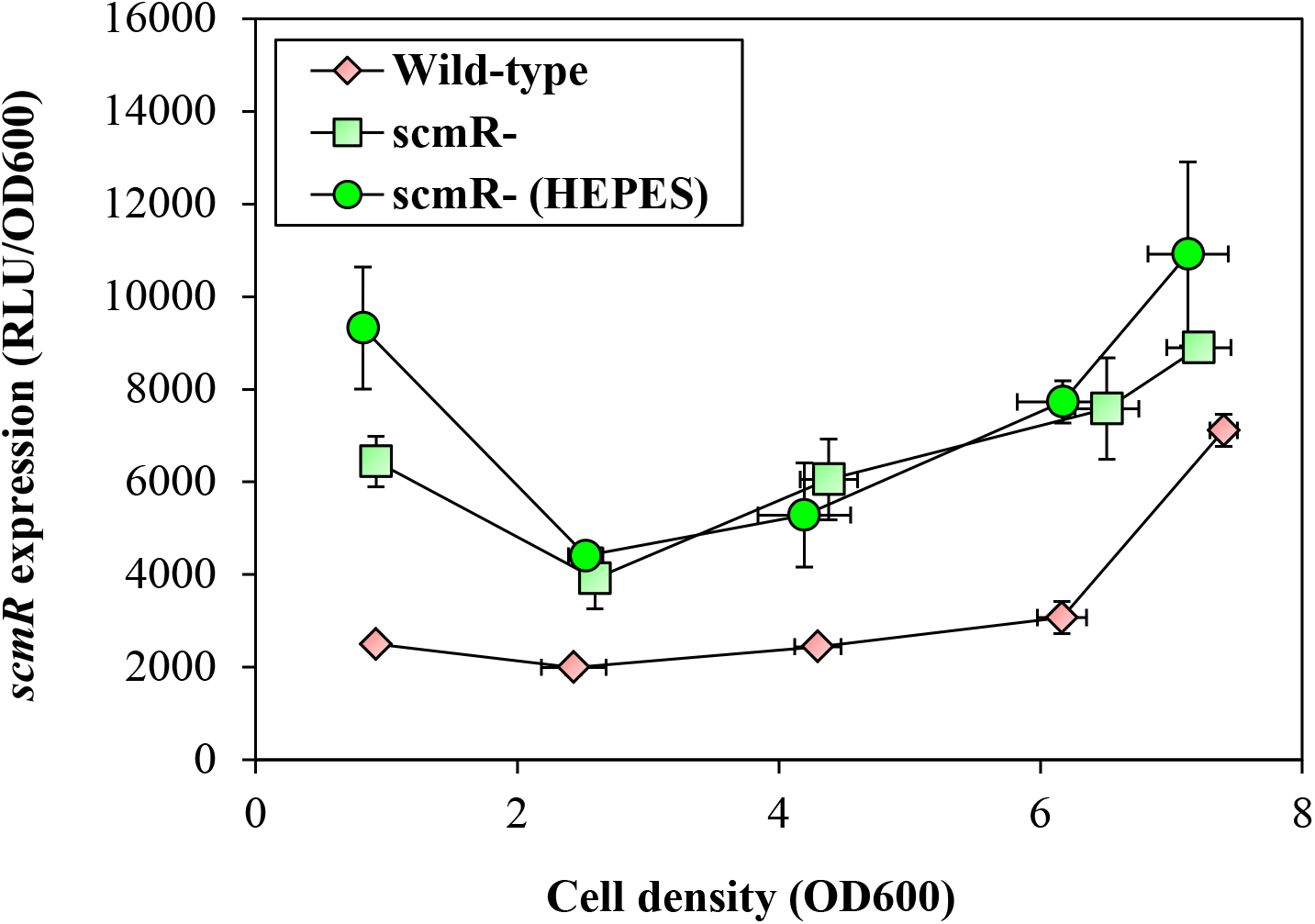
The *scmR* gene is negatively autoregulated. The luciferase activity of the chromosomal *scmR-lux* transcriptional fusion was monitored at various times during growth in cultures of the *B. thailandensis* E264 wild-type strain and the *scmR*-mutant strain. Cultures were buffered with 100 mM HEPES. Water only was added to the controls. The error bars represent the standard deviations of the averages for three replicates. The luminescence is expressed in relative light units per optical density of the culture (RLU/OD_600_).

A heterologous host *E. coli* expression reporter system was developed to examine the possibility of direct interaction of ScmR with the promoter region of the *scmR* gene. *E. coli* DH5α recombinant strains were generated containing the chromosomal *scmR-lux* transcriptional reporter and either pMLS7 or pMLS7-*scmR* for constitutive expression of the ScmR transcriptional regulator. In this systems, ScmR did not repress *scmR* transcription (data not shown), suggesting that ScmR does not directly repress its own expression or that additional unknown factor(s), which might be absent in the *E. coli* background, are required for *scmR* negative autoregulation. Indeed, LTTRs generally function in association with ligands to modulate the expression of genes (27).

### ScmR represses virulence in the fruit fly model *D. melanogaster*

The cytotoxin malleilactone was reported to contribute to virulence of both *B. thailandensis* E264 and *B. pseudomallei* Bp82 (15, 34). Interestingly, a Δ*scmR* mutant of *B. thailandensis* E264, which overproduces malleilactone, is more virulent toward the *C. elegans* nematode model host in comparison with the wild-type strain (26). Accordingly, we found that our *scmR*-mutant was significantly more virulent than the wild-type strain using the *D. melanogaster* host model (*P* < 0.001) (Fig. 10). However, according to our transcriptomic data and in contrast to Mao *et al.* (26) observations, ScmR had no impact on the transcription of the *mal* gene cluster (malleilactone biosynthesis) (15, 16), or on the expression of the orphan LuxR-type transcriptional regulator BtaR4 (MalR) which activates *mal* genes, indicating that ScmR might not regulate malleilactone production under our conditions (data not shown). Hence, the ScmR-dependent regulation of pathogenicity in *B. thailandensis* E264 is not exclusively mediated through control of the biosynthesis of malleilactone.

**Figure 10.**
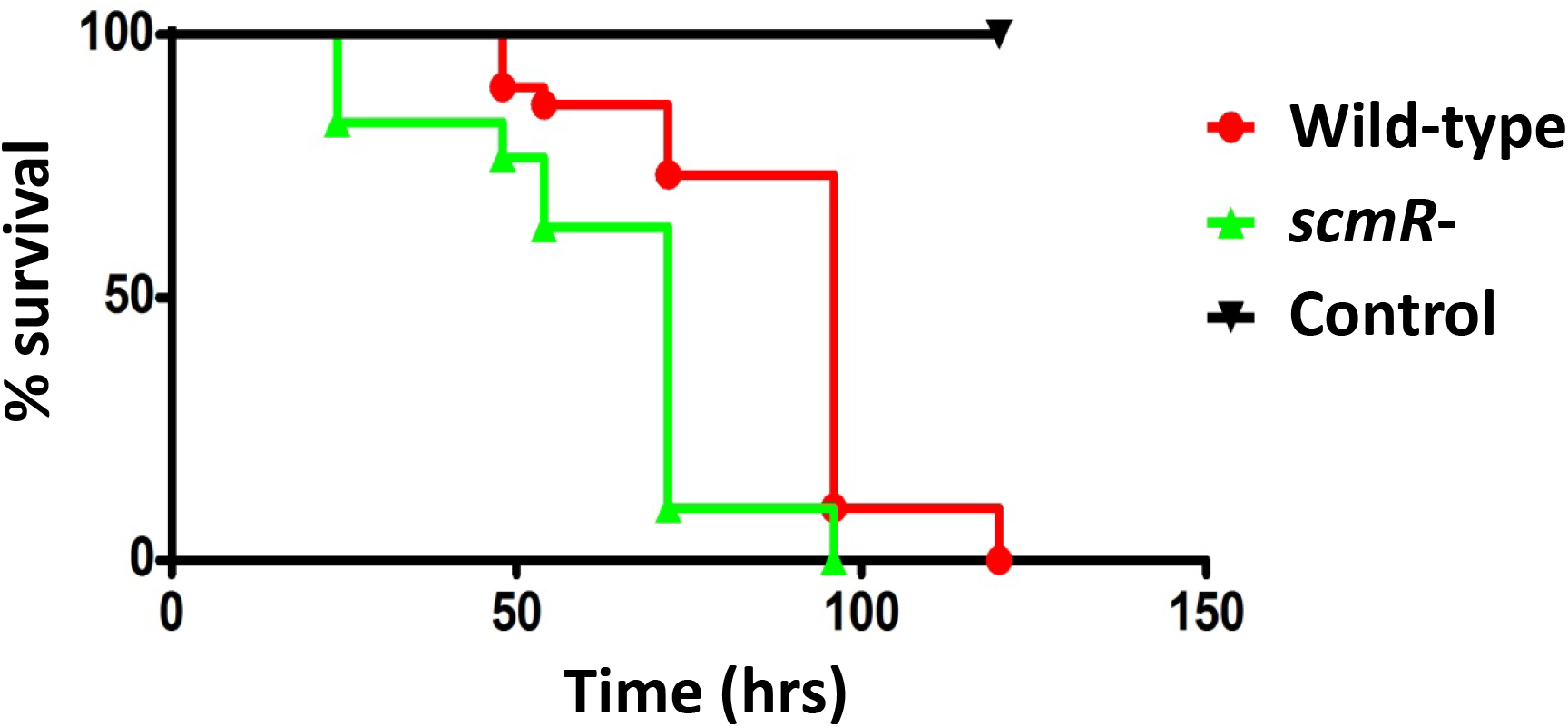
Virulence of the wild-type strain and of the *scmR*-mutant strain of *B. thailandensis* E264 toward the fruit fly *D. melanogaster*.

## Discussion

The function of the ScmR transcriptional regulator was recently addressed in *B. thailandensis*, revealing its importance in secondary metabolism regulation, as well as its involvement in the modulation of several QS-controlled phenotypes (26). While Mao *et al.* (26) defined the ScmR regulon during the stationary phase, we established the impact of ScmR on the expression of genes during the logarithmic phase of growth. It must be emphasized that the growth stage is an important variable when investigating QS, and this is especially relevant for *B. thailandensis*, a bacterium for which we reported significant differences in QS regulation depending on the growth stage (11). We confirmed that ScmR is a global regulator of gene expression in *B. thailandensis* E264 (Fig. 1A). Mao *et al.* (26) highlighted that ScmR modulates the production of the main AHL signaling molecules found in this bacterium, namely, C_8_-HSL, 3OHC_10_-HSL, and 3OHC_8_-HSL and we confirmed that AHLs biosynthesis is affected by ScmR as well (Fig. 3), which hints that ScmR might control the transcription of many genes through its effect on the QS-1, QS-2, and/or QS-3 systems. This is also further supported by the finding that ScmR modulates QS-controlled phenotypic traits, such as colony morphology, as well as pellicle and biofilm formation (26). Consistently, we noticed a considerable overlap between the ScmR-regulated genes and those controlled by QS (Fig. 2). Furthermore, we attested that the *scmR* gene is regulated by QS (Fig. 8), showing that ScmR is deeply integrated into the QS modulatory network of *B. thailandensis* E264. We assume that the QS-dependent regulation of *scmR* transcription allows tightly controlled coordination of the expression of genes.

Interestingly, we found that expression of many genes that encode transcriptional regulators, including the orphan QS transcriptional regulator BtaR5-encoding gene (Fig. 6E), is modulated either positively or negatively by ScmR (**Table S1**). Consequently, we propose that ScmR controls many genes through different and not mutually exclusive mechanisms: i.e. (i) regulation of AHL signaling molecules biosynthesis, (ii) direct binding of target genes, and (iii) indirect modulation of some genes via intermediate regulators. It will therefore be important to further investigate the molecular mechanism of action of ScmR to decipher between the directly and the indirectly ScmR-regulated genes. Moreover, the characterization of an ScmR-binding motif would contribute to the identification of promoters that are directly modulated versus those that are indirectly modulated.

The production of oxalic acid, which is required for pH homeostasis, is under QS control in several *Burkholderia* spp. (19, 31). Our RNA-Seq analyses confirmed the implication of AHLs in the regulation of expression of the oxalate biosynthetic gene *obc1*, and we showed that the transcription of *obc1* is stringently modulated by ScmR as well (Fig. 6A). Furthermore, we noticed that the impact of ScmR on *obc1* expression was more pronounced than the effect of AHLs (**Table S2**), suggesting that QS activates *obc1* transcription indirectly via positive regulation of the *scmR* gene. Whether the ScmR-dependent control of the *obc1* gene is direct or not remains to be determined. It is also possible that the ScmR-mediated control of the homeostasis of pH is not exclusively dependent on regulation of oxalate production. We indeed observed that several genes involved in ATP synthesis are modulated by ScmR (**Table S1**), as formerly noticed (26). Additionally, our RNA-Seq analyses revealed that ScmR stimulates expression of the putative lactate dehydrogenase LdhA-encoding gene, which is directly adjacent to *scmR*, and transcribed in the same direction in several *Burkholderia* spp. (26, 37) (**Fig. S2A**). Because lactate dehydrogenase, by reducing pyruvate to lactate, was suggested to affect pH (26), ScmR could also intervene in pH homeostasis through the activation of *ldhA* transcription. Indeed, Silva *et al.* (37) demonstrated that the *Burkholderia multivorans* ATCC 17616 homologue, called LdhR, influences pH homeostasis by activating both expression of the *ldhA* gene and lactate production. Still, we noticed no difference in pH between cultures of the *B. thailandensis* E264 wild-type strain and the *ldhA*-mutant (Fig. 3A), suggesting that LdhA is not involved in the ScmR-dependent control of pH homeostasis in *B. thailandensis* E264. Other discrepancies between the *B. thailandensis* E264 ScmR and the *B. multivorans* ATCC 17616 LdhR homologues were reported, such as their negative and positive effects on the development of biofilm, respectively (37). These findings indicate that these two proteins could be functionally different. More experiments will therefore be necessary to further understand the precise underlying molecular mechanism of action of ScmR in the control of pH homeostasis in *B. thailandensis*.

Since AHLs are hydrolyzed rapidly under alkaline pH conditions (32, 33), we propose that the impact of ScmR on the production of C_8_-HSL, 3OHC_10_-HSL, and 3OHC_8_-HSL might result, *inter alia*, from its influence on pH homeostasis (Fig. 7), This would then explain why no visible change in expression from the *btaI1*, *btaI2*, and *btaI3* promoters was observed in the *scmR*-mutant compared to the wild-type strain (data not shown). However, the ScmR-dependent regulation of the QS-1, QS-2, and/or QS-3 systems might be more complex and will need further investigation.

In agreement with the fact that LTTRs are typically negatively autoregulated (27), we found that ScmR represses its own expression (Fig. 9). Still, we saw no direct effect of the ScmR transcriptional regulator on the promoter region of the *scmR* gene when co-expressed together in a heterologous host. An explanation could be that *scmR* negative autoregulation requires additional modulatory elements, including molecular ligands. Indeed, ligands are recognized as being crucial for the function of LTTRs (27). They frequently participate to a self-inducing loop in which a product or an intermediate product of a given metabolic/synthesis pathway that is commonly stimulated by an LTTR acts as the ligand required for transcriptional regulation (27). Therefore, it will be important to determine the putative ligand(s) of ScmR in order to uncover the precise regulatory mechanism underlying *scmR* negative autoregulation.

While the pathogenicity of *B. multivorans* ATCC 17616 in the *Galleria mellonella* larvae model of infection is apparently not regulated by the ScmR homologue LdhR (37), the loss of ScmR in both *B. thailandensis* E264 and *B. pseudomallei* Bp82 resulted in hypervirulence toward the model host *C. elegans* (26, 34). In addition, Melanson *et al.* (38) demonstrated that the rice pathogen *B. glumae* 336gr-1 ScmR homologue, called NtpR, is a negative regulator of toxoflavin production, which is considered the primary virulence factor of *B. glumae*, suggesting that similarly to the *B. thailandensis* E264 and *B. pseudomallei* Bp82 ScmR homologues, NtpR influences virulence. In *B. thailandensis* and *B. pseudomallei*, ScmR was proposed to modulate the infection process by repressing the biosynthesis of malleilactone (26, 34). We demonstrated that ScmR of *B. thailandensis* E264 contributes to pathogenicity using the *D. melanogaster* host model (Fig. 10). However, we observed no difference in the transcription of the *mal* gene cluster, which encodes the principal enzymes responsible for malleilactone biosynthesis (15, 34), between the *B. thailandensis* E264 wild-type and the *scmR*-mutant strains, suggesting that ScmR is not necessarily involved in the production of the cytotoxin malleilactone. Any negative effect of ScmR on malleilactone biosynthesis *in vivo* is currently unknown. Still, we do not exclude the possibility that ScmR reduces virulence by modulating the expression of additional virulence/survival factors. For instance, we highlighted that expression of the *bsa* T3SS genes, which are crucial for the pathogenicity of both *B. pseudomallei* and *B. mallei* (39, 40), is repressed by ScmR (**Table S1**). The involvement of other potential virulence factors in the ScmR-mediated control of pathogenicity in *B. thailandensis* is currently under investigation.

## Materials and methods

### Bacterial strains and culture conditions

The bacterial strains used in this study are listed in **Table S3**. Unless otherwise stated, all bacteria were cultured at 37°C in tryptic soy broth (TSB; BD Difco, Mississauga, Ontario, Canada), with shaking (240 rpm) in a TC-7 roller drum (New Brunswick, Canada), or on Petri dishes containing TSB solidified with 1.5% agar. When required, antibiotics were used at the following concentrations: 200 μg/mL tetracycline (Tc), 100 μg/mL kanamycin (Km), and 100 μg/mL trimethoprim (Tp) for *B. thailandensis* E264, while Tc was used at 15 μg/mL for *E. coli* DH5α. All measurements of optical density at 600 nm (OD_600_) were acquired with a Thermo Fisher Scientific NanoDrop ND-1000 spectrophotometer.

### Construction of plasmids

Plasmids used in this study are described in **Table S4**. Amplification of the promoter region of *scmR* was performed from genomic DNA from *B. thailandensis* E264 using the appropriate primers (**Table S5**). The amplified product was digested with the FastDigest restriction enzymes *Xho*I and *Bam*HI (Thermo Fisher Scientific) and inserted by T4 DNA ligase (Bio Basic, Inc., Markham, ON, Canada) within the corresponding restriction sites in the mini-CTX-*lux* plasmid (41), generating the transcriptional reporter pSLG01. All primers were from Alpha DNA (Montreal, Quebec, Canada).

### Construction of reporter strains

Chromosomal integration of the mini-CTX-*scmR-lux* transcriptional reporter at the *attB* locus in *B. thailandensis* E264 strains was performed through conjugation with the auxotrophic *E. coli* χ7213, as described previously (12). Successful chromosomal insertion of *scmR*-*lux* was confirmed by PCR using appropriate primers.

### RNA isolation

Total RNA of *B. thailandensis* E264 cultures at an OD_600_ of 4.0 was extracted with the PureZOL RNA isolation reagent (Bio-Rad Laboratories, Mississauga, ON, Canada) and treated twice with the TURBO DNA-Free kit (Ambion Life Technologies, Inc., Burlington, ON, Canada) according to the manufacturer’s instructions. Extractions were done on two different bacterial cultures for RNA-Sequencing (RNA-Seq) analysis and on three different bacterial cultures for quantitative reverse transcription-PCR (qRT-PCR) and reverse transcription-PCR (RT-PCR) experiments. Quality and purity controls were confirmed by agarose gel electrophoresis and UV spectrophotometric analysis, respectively. Quantification of total RNA was accomplished on a Corbett Life Science Rotor-Gene 6000 thermal cycler using the QuantiFluor RNA system (Promega, Madison, WI, USA), according to the manufacturer’s protocol.

### RNA-Seq libraries construction and sequencing

The RNA-Seq libraries construction and sequencing using an Illumina HiSeq 2000 PE100 were performed by the McGill University and Génome Québec Innovation Centre (Montreal, QC, Canada). The RNA-Seq libraries were prepared using the TruSeq stranded mRNA sample preparation kit (Illumina, Inc., San Diego, CA, USA) and the Ribo-Zero rRNA removal kit (Epicentre, Madison, WI, USA).

### RNA-Seq mapping and analyses

All computations were made on the supercomputer Briarée from the Université de Montréal, managed by Calcul Québec and Compute Canada. Raw reads were filtered to remove low quality reads using the FASTX toolkit by discarding any reads with more than 10% nucleotides with a PHRED score < 20. Reads were then aligned with the reference genome (the GenBank accession no. for chromosome 1 of strain E264 is <u>CP000086.1</u> and for chromosome 2, it is <u>CP000085.1</u>) using Bowtie (v 2.2.3) with default parameters.

Chromosome 1 and chromosome 2 sequence alignments were separately processed to allow expression analysis between the two chromosomes. SAMtools (v 0.1.18) and BEDtools (v 2.20.1) were used for the generation of sam and bam files, respectively. The GC content of *B. thailandensis* E264 genes was calculated using BEDtools (v 2.20.1), prior to normalization. Normalization of the read count was done using the RPKM normalization function of the NOIseq package in R (42). To exclude features with low read counts, a low count filter was applied using a CPM method with a CPM value of 1 and a cutoff of 100 for the coefficient of variation. Cutoff values of 3-fold were used to consider differential expression biologically significant.

### LC-MS/MS quantification of AHLs

The concentrations of AHLs were determined from samples of *B. thailandensis* E264 cultures obtained at different time points during bacterial growth, by liquid chromatography coupled to tandem mass spectrometry (LC-MS/MS). The samples were prepared and analyzed as described previously (43). 5,6,7,8-Tetradeutero-4-hydroxy-2-heptylquinoline (HHQ-d4) was used as an internal standard. All experiments were performed in triplicate and conducted at least twice independently. For experiments with additions of 4-(2-hydroxyethyl)-1-piperazineethanesulfonic acid (HEPES), cultures were buffered or not with 100 mM HEPES (Sigma-Aldrich Co., Oakville, ON, Canada) from a stock prepared in ultrapure water. Water only was added to the controls.

### Quantitative reverse transcription-PCR and reverse transcription-PCR experiments

cDNA synthesis was performed using the iScript reverse transcription supermix (Bio-Rad Laboratories), and amplification was accomplished on a Corbett Life Science Rotor-Gene 6000 thermal cycler using the SsoAdvanced universal SYBR green supermix (Bio-Rad Laboratories), according to the manufacturer’s protocol. The reference gene was *ndh* (44). The *ndh* gene displayed stable expression under the different genetic contexts tested. All primers used for cDNA amplification are presented in **Table S6**. Differences in gene expression between *B. thailandensis* E264 strains were calculated using the 2^−ΔΔCT^ formula (45). A threshold of 0.5 was chosen as significant. All experiments were performed in triplicate and conducted at least twice independently.

### Measurement of the activity of *scmR*-*lux* reporter

Expression from the promoter region of *scmR* was quantified by measuring the luminescence of *B. thailandensis* E264 cultures carrying the corresponding chromosomal mini-CTX-*lux* transcriptional reporter, as described previously (11). Overnight bacterial cultures were diluted in TSB to an initial OD_600_ of 0.1 and incubated as indicated above. The luminescence was regularly determined from culture samples using a multi-mode microplate reader (Cytation 3; BioTek Instruments, Inc., Winooski, VT, USA) and expressed in relative light units per optical density of the culture (RLU/OD_600_). All experiments were performed with three biological replicates and repeated at least twice.

### Infection of D. melanogaster

The fruit flies were infected by feeding according to the previously described protocol (46). Briefly, 1 g of fruit fly dry medium was put into infection vials. Bacteria were harvested from LB-grown cultures adjusted to an OD_600_ of 4.0 by centrifugation at 10,000 x *g* for 5 min. The pellets were suspended in 0.02X PBS containing 1 mM CaCl_2_ and 1 mM MgCl_2_, as well as 500 µg/mL ampicillin (Ap) to avoid infection with nonspecific bacteria. Two mL of bacterial suspension were added to the dry food. Six-seven days-old male flies were anesthetized with CO_2_ and added to the vials by group of 10. The control vials contained the PBS solution only. Fly survival was scored daily and survival curves were processed with GraphPad Prism 5 (GraphPad Software, Inc., San Diego, CA, USA) to perform a statistical log-rank (Mantel-Cox) test.

### Data analysis

Unless stated otherwise, data are reported as means ± standard deviations (SD). Statistical analyses were performed with the R software version 3.3.3 (http://www.R-project.org/) using one-way analysis of variance (ANOVA). Probability values of less than 0.05 were considered significant.

## Funding information

This study was supported by Canadian Institutes of Health Research (CIHR) operating grants MOP-97888 and MOP-142466 to ED. ED holds the Canada Research Chair in Sociomicrobiology. The funders had no role in study design, data collection and interpretation, or the decision to submit the work for publication.

## Acknowledgments

We thank Everett Peter Greenberg (Department of Microbiology, University of Washington School of Medecine, Seattle, WA, USA) for providing the *B. thailandensis* E264 strains. We especially thank Sylvain Milot, François D’Heygere, and Koyomi Ozaki for their technical help.

## References

1. Fuqua WC, Winans SC, Greenberg EP. 1994. Quorum sensing in bacteria: the LuxR-LuxI family of cell density-responsive transcriptional regulators. J Bacteriol 176:269–75.

2. Nealson KH, Platt T, Hastings JW. 1970. Cellular control of the synthesis and activity of the bacterial luminescent system. J Bacteriol 104:313–22.

3. Fuqua WC, Greenberg EP. 2002. Listening in on bacteria: acyl-homoserine lactone signalling. Nat Rev Mol Cell Biol 3:685–95.

4. Majerczyk CD, Greenberg EP, Chandler JR. 2013. Quorum Sensing in *Burkholderia*. In Vasil, M, and Darwin, A (ed), Regulation of Bacterial Virulence ASM Press, Washington, DC doi:10.1128/9781555818524.ch3:p 40-57.

5. Brett PJ, DeShazer D, Woods DE. 1998. *Burkholderia thailandensis* sp. nov., a *Burkholderia pseudomallei*-like species. Int J Syst Bacteriol 48 Pt 1:317–20.

6. Haraga A, West TE, Brittnacher MJ, Skerrett SJ, Miller SI. 2008. *Burkholderia thailandensis* as a model system for the study of the virulence-associated type III secretion system of *Burkholderia pseudomallei*. Infect Immun 76:5402–11.

7. Ulrich RL, Deshazer D, Brueggemann EE, Hines HB, Oyston PC, Jeddeloh JA. 2004. Role of quorum sensing in the pathogenicity of *Burkholderia pseudomallei*. J Med Microbiol 53:1053–64.

8. Ulrich RL, Deshazer D, Hines HB, Jeddeloh JA. 2004. Quorum sensing: a transcriptional regulatory system involved in the pathogenicity of *Burkholderia mallei*. Infect Immun 72:6589–96.

9. Ulrich RL, Hines HB, Parthasarathy N, Jeddeloh JA. 2004. Mutational analysis and biochemical characterization of the *Burkholderia thailandensis* DW503 quorum-sensing network. J Bacteriol 186:4350–60.

10. Majerczyk CD, Brittnacher M, Jacobs M, Armour CD, Radey M, Schneider E, Phattarasokul S, Bunt R, Greenberg EP. 2014. Global analysis of the *Burkholderia thailandensis* quorum sensing-controlled regulon. J Bacteriol 196:1412–24.

11. Le Guillouzer S, Groleau MC, Déziel E. 2017. The Complex Quorum Sensing Circuitry of *Burkholderia thailandensis* Is Both Hierarchically and Homeostatically Organized. mBio 8:e01861–17.

12. Le Guillouzer S, Groleau MC, Déziel E. 2018. Two *rsaM* homologues encode central regulatory elements modulating quorum sensing in *Burkholderia thailandensis*. J Bacteriol 200:e00727–17.

13. Chandler JR, Duerkop BA, Hinz A, West TE, Herman JP, Churchill ME, Skerrett SJ, Greenberg EP. 2009. Mutational analysis of *Burkholderia thailandensis* quorum sensing and self-aggregation. J Bacteriol 191:5901–9.

14. Duerkop BA, Varga J, Chandler JR, Peterson SB, Herman JP, Churchill ME, Parsek MR, Nierman WC, Greenberg EP. 2009. Quorum-sensing control of antibiotic synthesis in *Burkholderia thailandensis*. J Bacteriol 191:3909–18.

15. Biggins JB, Ternei MA, Brady SF. 2012. Malleilactone, a polyketide synthase-derived virulence factor encoded by the cryptic secondary metabolome of *Burkholderia pseudomallei* group pathogens. J Am Chem Soc 134:13192–5.

16. Truong TT, Seyedsayamdost M, Greenberg EP, Chandler JR. 2015. A *Burkholderia thailandensis* Acyl-Homoserine Lactone-Independent Orphan LuxR Homolog That Activates Production of the Cytotoxin Malleilactone. J Bacteriol 197:3456–62.

17. Valade E, Thibault FM, Gauthier YP, Palencia M, Popoff MY, Vidal DR. 2004. The PmlI-PmlR quorum-sensing system in *Burkholderia pseudomallei* plays a key role in virulence and modulates production of the MprA protease. J Bacteriol 186:2288–94.

18. Song Y, Xie C, Ong YM, Gan YH, Chua KL. 2005. The BpsIR quorum-sensing system of *Burkholderia pseudomallei*. J Bacteriol 187:785–90.

19. Goo E, Majerczyk C, An JH, Chandler JR, Seo YS, Ham H, Lim JY, Kim H, Lee B, Jang MS, Greenberg EP, Hwang I. 2012. Bacterial quorum sensing, cooperativity, and anticipation of stationary-phase stress. Proc Natl Acad Sci U S A 109:19775–80.

20. Majerczyk CD, Brittnacher MJ, Jacobs MA, Armour CD, Radey MC, Bunt R, Hayden HS, Bydalek R, Greenberg EP. 2014. Cross-species comparison of the *Burkholderia pseudomallei*, *Burkholderia thailandensis*, and *Burkholderia mallei* quorum-sensing regulons. J Bacteriol 196:3862–71.

21. Tseng BS, Majerczyk CD, Passos da Silva D, Chandler JR, Greenberg EP, Parsek MR. 2016. Quorum Sensing Influences *Burkholderia thailandensis* Biofilm Development and Matrix Production. J Bacteriol 198:2643–50.

22. Ulrich RL. 2004. Quorum quenching: enzymatic disruption of *N*-acylhomoserine lactone-mediated bacterial communication in *Burkholderia thailandensis*. Appl Environ Microbiol 70:6173–80.

23. Ooi WF, Ong C, Nandi T, Kreisberg JF, Chua HH, Sun G, Chen Y, Mueller C, Conejero L, Eshaghi M, Ang RM, Liu J, Sobral BW, Korbsrisate S, Gan YH, Titball RW, Bancroft GJ, Valade E, Tan P. 2013. The condition-dependent transcriptional landscape of *Burkholderia pseudomallei*. PLoS Genet 9:e1003795.

24. Ramli NS, Eng Guan C, Nathan S, Vadivelu J. 2012. The effect of environmental conditions on biofilm formation of *Burkholderia pseudomallei* clinical isolates. PLoS One 7:e44104.

25. Mongkolrob R, Taweechaisupapong S, Tungpradabkul S. 2015. Correlation between biofilm production, antibiotic susceptibility and exopolysaccharide composition in *Burkholderia pseudomallei bpsI*, *ppk*, and *rpoS* mutant strains. Microbiol Immunol 59:653–63.

26. Mao D, Bushin LB, Moon K, Wu Y, Seyedsayamdost MR. 2017. Discovery of *scmR* as a global regulator of secondary metabolism and virulence in *Burkholderia thailandensis* E264. Proc Natl Acad Sci U S A 114:E2920–E2928.

27. Maddocks SE, Oyston PC. 2008. Structure and function of the LysR-type transcriptional regulator (LTTR) family proteins. Microbiology 154:3609–23.

28. Vial L, Lépine F, Milot S, Groleau MC, Dekimpe V, Woods DE, Déziel E. 2008. *Burkholderia pseudomallei*, *B. thailandensis*, and *B. ambifaria* produce 4-hydroxy-2-alkylquinoline analogues with a methyl group at the 3 position that is required for quorum-sensing regulation. J Bacteriol 190:5339-52.

29. Biggins JB, Gleber CD, Brady SF. 2011. Acyldepsipeptide HDAC inhibitor production induced in *Burkholderia thailandensis*. Org Lett 13:1536–9.

30. Majerczyk CD, Schneider E, Greenberg EP. 2016. Quorum sensing control of Type VI secretion factors restricts the proliferation of quorum-sensing mutants. Elife 5.

31. Goo E, An JH, Kang Y, Hwang I. 2015. Control of bacterial metabolism by quorum sensing. Trends Microbiol 23:567–76.

32. Byers JT, Lucas C, Salmond GP, Welch M. 2002. Nonenzymatic turnover of an *Erwinia carotovora* quorum-sensing signaling molecule. J Bacteriol 184:1163–71.

33. Yates EA, Philipp B, Buckley C, Atkinson S, Chhabra SR, Sockett RE, Goldner M, Dessaux Y, Camara M, Smith H, Williams P. 2002. *N-*acylhomoserine lactones undergo lactonolysis in a pH-, temperature-, and acyl chain length-dependent manner during growth of *Yersinia pseudotuberculosis* and *Pseudomonas aeruginosa*. Infect Immun 70:5635–46.

34. Klaus JR, Deay J, Neuenswander B, Hursh W, Gao Z, Bouddhara T, Williams TD, Douglas J, Monize K, Martins P, Majerczyk C, Seyedsayamdost MR, Peterson BR, Rivera M, Chandler JR. 2018. Malleilactone Is a *Burkholderia pseudomallei* Virulence Factor Regulated by Antibiotics and Quorum Sensing. J Bacteriol 200.

35. O’Grady EP, Viteri DF, Malott RJ, Sokol PA. 2009. Reciprocal regulation by the CepIR and CciIR quorum sensing systems in *Burkholderia cenocepacia*. BMC Genomics 10:441.

36. Chambers CE, Lutter EI, Visser MB, Law PP, Sokol PA. 2006. Identification of potential CepR regulated genes using a cep box motif-based search of the *Burkholderia cenocepacia* genome. BMC Microbiol 6:104.

37. Silva IN, Ramires MJ, Azevedo LA, Guerreiro AR, Tavares AC, Becker JD, Moreira LM. 2017. Regulator LdhR and d-Lactate Dehydrogenase LdhA of *Burkholderia multivorans* Play Roles in Carbon Overflow and in Planktonic Cellular Aggregate Formation. Appl Environ Microbiol 83.

38. Melanson RA, Barphagha I, Osti S, Lelis TP, Karki HS, Chen R, Shrestha BK, Ham JH. 2017. Identification of new regulatory genes involved in the pathogenic functions of the rice-pathogenic bacterium *Burkholderia glumae*. Microbiology 163:266–279.

39. Warawa J, Woods DE. 2005. Type III secretion system cluster 3 is required for maximal virulence of *Burkholderia pseudomallei* in a hamster infection model. FEMS Microbiol Lett 242:101–8.

40. Ulrich RL, DeShazer D. 2004. Type III secretion: a virulence factor delivery system essential for the pathogenicity of *Burkholderia mallei*. Infect Immun 72:1150–4.

41. Becher A, Schweizer HP. 2000. Integration-proficient *Pseudomonas aeruginosa* vectors for isolation of single-copy chromosomal *lacZ* and *lux* gene fusions. Biotechniques 29:948–50, 952.

42. Tarazona S, Garcia-Alcalde F, Dopazo J, Ferrer A, Conesa A. 2011. Differential expression in RNA-seq: a matter of depth. Genome Res 21:2213–23.

43. Chapalain A, Groleau MC, Le Guillouzer S, Miomandre A, Vial L, Milot S, Déziel E. 2017. Interplay between 4-Hydroxy-3-Methyl-2-Alkylquinoline and *N*-Acyl-Homoserine Lactone Signaling in a *Burkholderia cepacia* Complex Clinical Strain. Front Microbiol 8:1021.

44. Subsin B, Chambers CE, Visser MB, Sokol PA. 2007. Identification of genes regulated by the *cepIR* quorum-sensing system in *Burkholderia cenocepacia* by high-throughput screening of a random promoter library. J Bacteriol 189:968–79.

45. Livak KJ, Schmittgen TD. 2001. Analysis of relative gene expression data using real-time quantitative PCR and the 2(-Delta Delta C(T)) Method. Methods 25:402–8.

46. Pilatova M, Dionne MS. 2012. *Burkholderia thailandensis* is virulent in *Drosophila melanogaster*. PLoS One 7:e49745.

