## Supplemental Figures S1-S3 and Tables S3-S6 for "ScmR, a global regulator of gene expression, quorum sensing, pH homeostasis, and virulence in *Burkholderia thailandensis*"

| | | | |
|---|---|---|---|
| *Vibrio fischeri* | *luxI* | (−32 to −13) | ACCTGTAGGATCGTACAGGT |
| *Burkholderia mallei* | *scmR* | (−82 to −63) | CCAGTGCTGATCCGACAGCC |
| *Pseudomonas aeruginosa* | *rhlI* | (−155 to −136) | CCCTACCAGATCTGGCAGGT |
| *Burkholderia thailandensis* | *rsaM1* | (−54 to −35) | CGCTGTCATACTTGCTAGGT |
| *Burkholderia pseudomallei* | *scmR* | (−82 to −63) | CCAGTGCTGATCCGACAGCC |
| *Pseudomonas aeruginosa* | *lasI* | (−75 to −56) | ATCTATCTCATTTGCTAGTT |
| *Burkholderia cenocepacia* | *cepI* | (−82 to −63) | CCCTGTAAGAGTTACCAGTT |

**Figure S1. The promoter region of the ScmR-encoding gene contains a putative *lux* box sequence.** The *scmR* gene encoding ScmR possesses in its promoter region a putative *lux* box sequence, which is homologous to characterized *lux* box sequences in Proteobacteria. The start and stop positions of each sequence are indicated in parentheses relative to the translational start site.

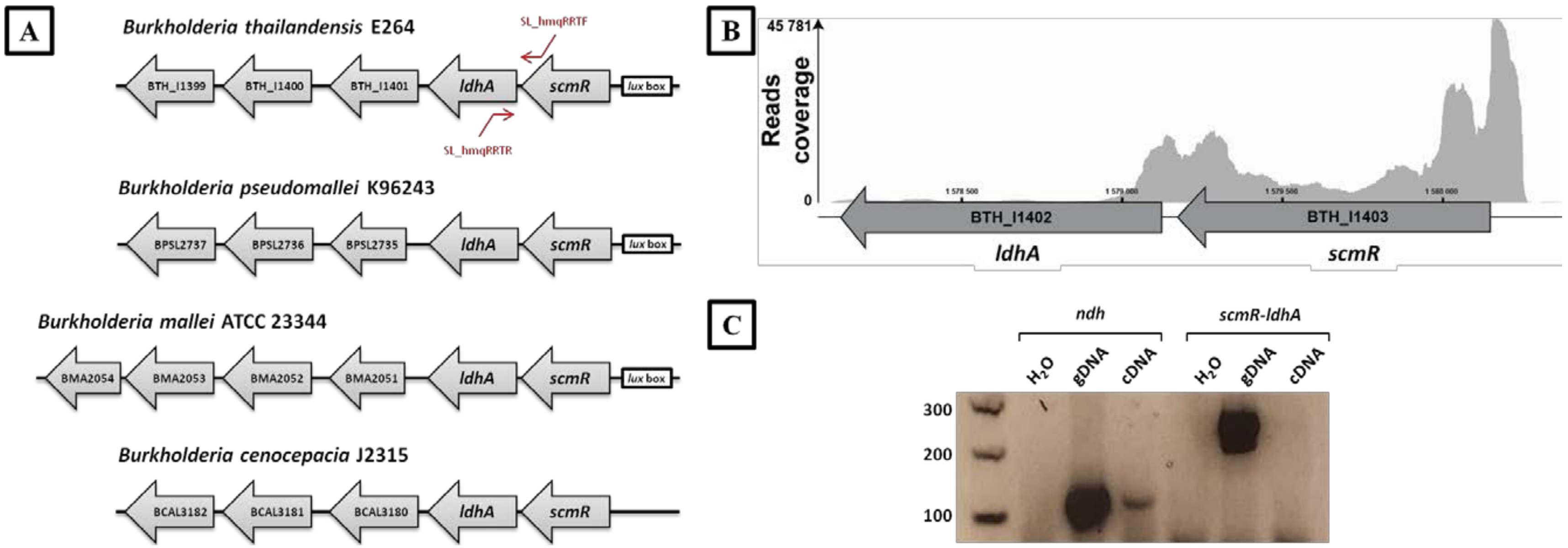


**Figure S2. Examination of the genetic organization of *scmR*.** (A) The *scmR* gene is directly adjacent to *ldhA* on the genome of *B. thailandensis* E264. The SL_hmqRRTF and SL_hmqRRTR primer pair was used to amplify the intergenic region of the *scmR* and *ldhA* genes. The promoter region of *scmR* contains a putative *lux* box sequence as reported formerly (Mao *et al.*, 2017). *B. pseudomallei* K96243, *B. mallei* ATCC 23344, and *B. cenocepacia* J2315 possess an *scmR* homologue. A putative *lux* box sequence is present in the promoter region of the *B. pseudomallei* K96243 and *B. mallei* ATCC 23344 ScmR-encoding gene (CCAGTGCTGATCCGACAGCC), but not in the promoter region of the *B. cenocepacia* J2315 ScmR-encoding gene as reported formerly (1). (B) Transcriptomic analyses obtained by RNA-Seq showing the genetic arrangement of the ScmR-encoding gene of *B. thailandensis* E264. (C) RT-PCR experiments were performed to examine cotranscription of the *scmR* gene with *ldhA* and analyzed by agarose gel electrophoresis. The notation *ndh* indicates that the PCR reactions were conducted using the SLG_qRT-PCR_ndh_F and SLG_qRT-PCR_ndh_R primer pair with $H_2O$ as a negative control, genomic DNA (gDNA) as a positive control, or complementary DNA (cDNA) of *B. thailandensis* E264. These reactions testify to the efficiency of the RT-PCR experiments as revealed by the presence of a PCR amplification product in the cDNA lane. The notation *scmR-ldhA* indicates that the PCR reactions were carried out with the SL_hmqRRTF and SL_hmqRRTR primer pair. The absence of PCR amplification product in the cDNA lane reveals that *scmR* is not cotranscribed with the *ldhA* gene. The molecular marker is the DNA Ladder 100 bp (New England Biolabs, Inc., Whitby, ON, Canada).

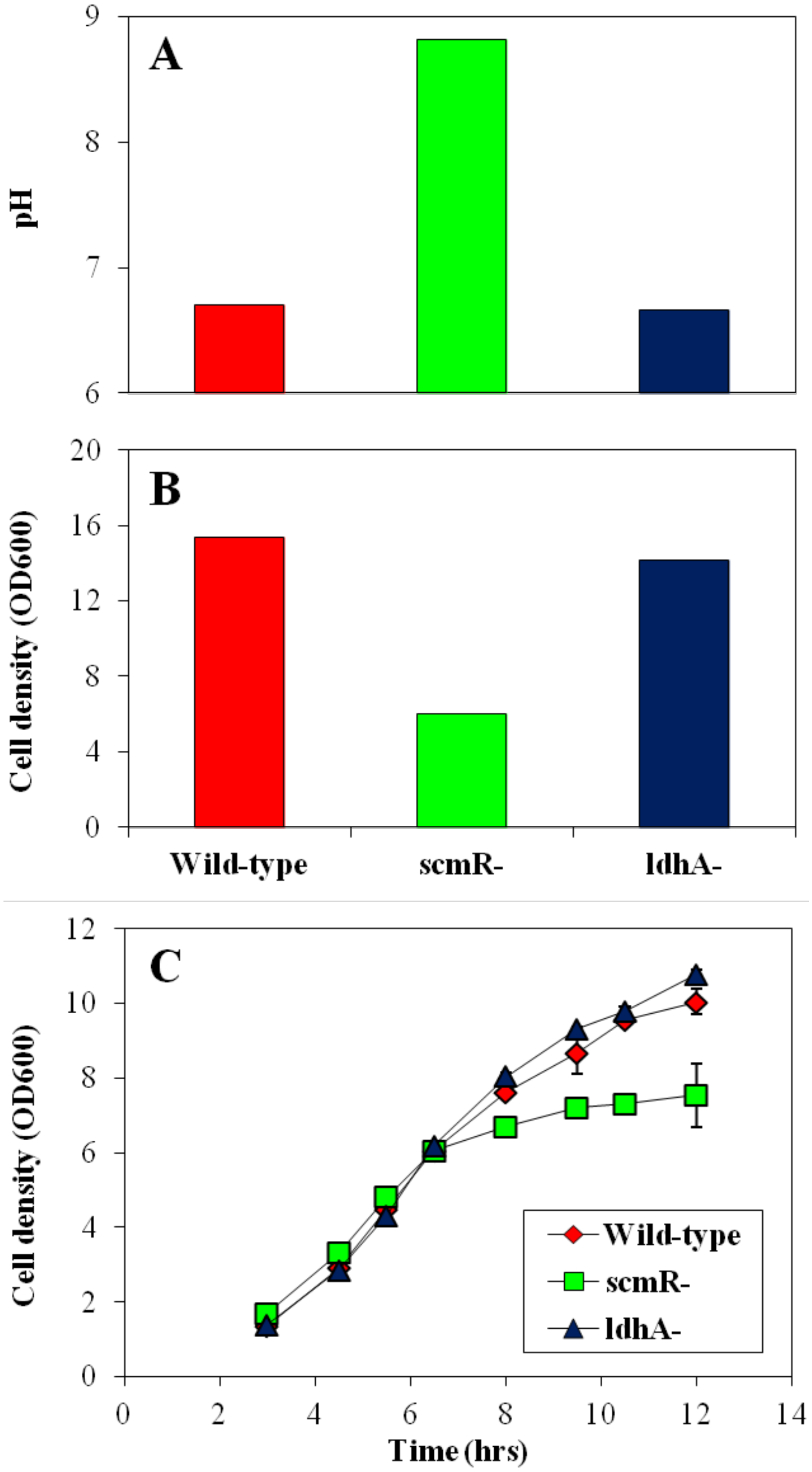


**Figure S3. pH homeostasis is not affected by the *B. thailandensis* E264 LdhA lactate dehydrogenase.** (A) pH value was measured with a pH electrode and meter (Mettler-Toledo, Mississauga, ON, Canada), during the stationary growth phase in cultures of the *B. thailandensis* E264 wild-type strain and the *scmR*- and *ldhA*- mutant strains. Cultures were buffered with 100 mM HEPES. Water only was added to the controls. (B) The cell density was monitored by measuring turbidity, expressed in 600 nm absorption units ($OD_{600}$). (C) The *B. thailandensis* E264 wild-type strain and the *scmR*- and *ldhA*- mutant strains growth curves. The error bars represent the standard deviations of the averages for three replicates.

**Tables S1 and S2 : see Excel file.**

**Table S3. Bacterial strains used in this study.**

| Strains | Description | Reference |
|---|---|---|
| ***E. coli*** | | |
| **χ7213** | *thr*-1, *leuB*6, *fhuA*21, *lacY*1, *glnV*44, *recA*1, Δ*asdA*4, Δ(*zhf*-2::Tn*10*), *thi*-1, RP4-2-Tc :: Mu [λ *pir*] | Lab collection |
| **DH5α** | F-, φ80d*lacZ*ΔM15, Δ(*lacZYA-argF*)U169, *deoR*, *recA*1, *endA*1, *hsdR*17(rk-, mk+), *phoA*, *supE*44, λ-, *thi*-1, *gyrA*96, *relA*1 | Lab collection |
| ***B. thailandensis*** | | |
| **E264** | Wild-type | (2) |
| **ED1023** | E264 *scmR*::pUT-mini-Tn*5*-Km; $Km^R$ | (Le Guillouzer *et al.,* unpublished) |
| **JBT112** | E264 Δ*btaI1* Δ*btaI2* Δ*btaI3* | (3) |
| **JBT107** | E264 Δ*btaR1* | (3) |
| **JBT108** | E264 Δ*btaR2* | (3) |
| **JBT109** | E264 Δ*btaR3* | (3) |
| **BT08944** | E264 *ldhA*::Is*lacZ*-PrhaBo-Tp/FRT;$Tp^R$ | (4) |
| **ED351**5 | E264::*scmR-lux* | This study |
| **ED351**7 | E264 *scmR*-::*scmR-lux* | This study |

**Table S4. Plasmids used in this study.**

| Plasmids | Description | Source |
|---|---|---|
| **mini-CTX-*lux*** | Integration vector with promoterless *luxCDABE*; $Tc_R$ | (5) |
| **pSLG01** | *scmR* promoter inserted in *Xho*I-*Bam*HI restriction sites in mini-CTX-*lux*; $Tc^R$ | This study |

**Table S5. Primers used for PCR.**

| Genes | Oligonucleotides | Sequences (5’ to 3’) |
|---|---|---|
| ***scmR*** | PhmqRF | CGCCTCGAGGAGTGATCGGTCGTGGACAT |
| | PhmqRR | CGCGGATCCTCGATACGCCGAGTTTCC |

**Table S6. Primers used for qRT-PCR and RT-PCR.**

| Genes | Oligonucleotides | Sequences (5' to 3') |
|---|---|---|
| ***ndh*** | SLG_qRT-PCR_ndh_F | ACCAGGGCGAATTGATCTC |
| | SLG_qRT-PCR_ndh_R | GATGACGAGCGTGTCGTATT |
| ***obc1*** | SLG_BTH_II1071_F | CTATCGTCGGTCGATCTGGT |
| | SLG_BTH_II1071_R | GTAGGTGTGGATCCGGTCGT |
| **BTH_I3204** | SLG_BTH_I3204_F | AAGCAGAACGGCCTGACTAA |
| | SLG_BTH_I3204_R | TGCTATCGAGCACCGTGTAG |
| ***bsaN*** | SLG_BTH_II0827_F | GAAATCGCGAAACTGGATGT |
| | SLG_BTH_II0827_R | TAACGGCACGTCATGAAAAC |
| **BTH_II0639** | SLG_BTH_II0639_F | GACTTTCCTCTCGCTTGCAG |
| | SLG_BTH_II0639_R | GCACGAGGATGATCGGATAC |
| ***btaR5*** | SLG_BTH_I1817_F | AACTTTCCCGAGCAATGGA |
| | SLG_BTH_I1817_R | GTCTCGTTCAGGACGACAGG |
| **BTH_II1209** | SLG_BTH_II1209_F | GTCCCTTCCCATTCACACTG |
| | SLG_BTH_II1209_R | TTAACGGCTTTGCCTAATGC |
| ***scmR*** | SLG_qRThmqR_F | CTTCGTATGTGTTGCCGAAC |
| | SLG_qRThmqR_R | ATGAGACGCGTGTTCAGATG |
| ***scmR-ldhA*** | SL_hmqRRTF | ACGCAGTTGTCCATCGTCTA |
| | SL_hmqRRTR | CGGCTGCTGAAAAGAATCAC |

### References


1. Chambers CE, Lutter EI, Visser MB, Law PP, Sokol PA. 2006. Identification of potential CepR regulated genes using a cep box motif-based search of the *Burkholderia cenocepacia* genome. BMC Microbiol 6:104.
2. Brett PJ, DeShazer D, Woods DE. 1998. *Burkholderia thailandensis* sp. nov., a *Burkholderia pseudomallei*-like species. Int J Syst Bacteriol 48 Pt 1:317-20.
3. Chandler JR, Duerkop BA, Hinz A, West TE, Herman JP, Churchill ME, Skerrett SJ, Greenberg EP. 2009. Mutational analysis of *Burkholderia thailandensis* quorum sensing and self-aggregation. J Bacteriol 191:5901-9.
4. Gallagher LA, Ramage E, Patrapuvich R, Weiss E, Brittnacher M, Manoil C. 2013. Sequence-defined transposon mutant library of *Burkholderia thailandensis*. MBio 4:e00604-13.
5. Becher A, Schweizer HP. 2000. Integration-proficient *Pseudomonas aeruginosa* vectors for isolation of single-copy chromosomal *lacZ* and *lux* gene fusions. Biotechniques 29:948-50, 952.
